# Proteomic stress response by a novel methanogen enriched from the Great Salt Lake

**DOI:** 10.64898/2026.01.29.702513

**Authors:** William C Christian, Zackary J Jay, Nikola Tolic, Carrie D Nicora, Reece Livingstone, Stavros Trimmer, Timothy R McDermott, Roland Hatzenpichler

**Author notes:** corresponding author: R.H.

## Abstract

Methanogenic archaea affect the climate through their production of the greenhouse gas, methane. However, it is unclear how a changing climate and other anthropogenic influences impact methanogen physiology and consequent methane flux. The Great Salt Lake (GSL) is an environment that has been heavily impacted by human activity; more than doubling its salt concentration since the last methanogen was cultured from it in 1985. In this study, we enriched a novel methanogen, for which we propose the name *Candidatus* Methanohalophilus hillemani, from the GSL at a time when its salinity reached a historical high. Interestingly, *Ca.* M. hillemani does not increase expression of energy-conservation or osmo-tolerance proteins when challenged with salinity or oxygen. In contrast, *Ca.* M. hillemani prioritizes trace metal uptake and immune functions in response to the presence of the sulfate-reducing bacterium *Desulfovermiculus*. 16S rRNA gene amplicon data from GSL shore soils with extremely high and variable methane flux indicated the presence of *Ca.* M. hillemani. Our results show that *Ca.* M. hillemani is active when challenged with environmental stressors and contributes to the methane flux emanating from the GSL.

**Importance:** Methanogens are microbes that affect the climate through their production of the greenhouse gas, methane. Changes in climate and land-use patterns are drying up saline lakes, damaging their unique economic and ecological value. As lake levels across the globe fall, it is unclear how methanogens and the amount of methane they produce will concurrently shift. In this study, we measured high methane output from the Great Salt Lake (GSL) across seasons and identified a novel methanogen as part of a larger methanogenic community that is responsible for these methane emissions. We cultured this novel methanogen from GSL sediments and determined that its methane production was largely unaffected by stress conditions. Our findings indicate that methanogens in saline environments, including a novel cultivated species, may be important sources of methane and will continue to produce methane as salinity increases.

## Introduction

Methane is a potent greenhouse gas that has contributed 0.3-0.8 °C to global warming since the start of the industrial revolution [1, 2]. Approximately 60% of methane emitted to the atmosphere is produced by methanogens, archaea that couple the production of methane to energy conservation [2, 3]. Due to the oxygen sensitivity of key enzymes in their metabolism, as well as the comparatively low free-energy yield of methanogenesis, it was long assumed that methanogens are restricted to anoxic environments with large stores of organic carbon. Specifically, wetlands, melting permafrosts, rice paddies, and animal digestive tracts are known for their high methane fluxes [2] and have been extensively studied, together with methanogens therein, due to their influence on climate change. However, many additional ecosystems contribute to the methane cycle. Environments experiencing fast changes due to anthropogenic influence are of particular interest because the magnitude and direction (i.e. emission vs. uptake) of methane flux can change on annual to decadal scales [4–7].

The Great Salt Lake (GSL) and other major terminal lakes around the globe are being altered by human activity [8–10]. Increased upstream water consumption and evaporation due to higher temperatures are causing lake levels to plummet [11–13]. In terminal lakes, a decrease in water volume is accompanied by an increase in salinity. It is well known that halophilic methanogens produce methane under hypersaline conditions [14], however, how they respond to increasing salinity, and how this impacts methane flux, is not well known. Recent studies suggest that increasing salinity in freshwater systems decreases methane emissions [15], but also that droughts will increase methane emissions from once-inundated sediments [16]. Studies specifically examining salinity gradients have shown mixed results, with some reporting no correlation between salinity and methane emissions [17] and others finding a negative relationship [18, 19]. These results have contradictory implications for methane emissions from drying terminal lakes, including the GSL. Unfortunately, data examining emissions from these unique environments is rare and to the best of our knowledge, only one study has quantified methane flux from a terminal lake [20], which measured low and spatially homogenous methane emissions from the GSL. Methane production has been observed from GSL sediments [21], and a halophilic methanogen, *Methanohalophilus mahii*, was isolated from the GSL in 1985 [22–24]. Despite these observations, many microbial community analyses of the GSL did not find or mention methanogens [25–27].

*M. mahii* is a methyl-dismutating methanogen that thrives at 7-12% NaCl [23]. The *Methanohalophilus* genus includes six described species and several strains [28, 29]. Besides the GSL, these species were isolated from deep sea salt brines, deep shale reservoirs, hypersaline bays, and salt pans [29–33]. Known members of the *Methanohalophilus* genus all share methyl-dismutating methanogenesis from MMA, DMA, TMA (with one exception [34]) and methanol, but are incapable of growing via acetoclastic or CO_2_-reducing hydrogenotrophic methanogenesis. The salinity range varies by organism, but they have been shown to grow with NaCl concentrations from 1% to 20% [28]. They tolerate osmotic stress through both ‘salt-in’ strategies with many ion symporters and antiporters, and via the biosynthesis of the compatible solute, glycine betaine [34]. All members of this genus also universally encode genes for the detoxification of O_2_, including thioredoxins, peroxiredoxins, alkylhydroperoxidases, and superoxide reductase [28].

In this study, we cultured a novel species of the genus *Methanohalophilus* from the GSL for which we propose the name *Candidatus* Methanohalophilus hillemani. Using metagenomics and metaproteomics, we describe its unique features and explore how this organism responds to environmental stressors, including salinity. Finally, we quantify methane flux from the GSL and identify the microbes responsible. We combine environmental, microbiological, and meta-omics techniques to examine how this methanogen responds to changes experienced in the GSL.

## Materials and methods

### Collection of samples and cultivation

Microbialite material was collected in July of 2022 from Bridger Bay in the South arm of the GSL (Supplementary Figures 1, 2). Methanogens were enriched by adding 10 mM MMA to 0.2 µm filter-sterilized GSL water. Lake water was eventually replaced by media designed by Paterek and Smith [22], originally used to isolate *Methanohalophilus mahii*, but adjusted to the salinity of the current GSL (16%).

*Methanohalophilus* enrichment cultures were grown in Paterek and Smith 1985 media [22], with differing concentrations of salts and TMA, and without the addition of volatile fatty acid solution and sodium sulfide. The headspace consisted of a 50/45/5 ratio of H_2_/N_2_/CO_2_ at ambient pressure (∼90 kPa in Bozeman, MT). The anoxic media included the following salts to reflect the salinity of GSL water from April 13^th^ 2023: 100 g/L NaCl, 45 g/L MgCl_2_, 5.5 g/L KCl, 1.5 g/L CaCl_2_, 0.13 g/L LiCl, 19 g/L Na_2_SO_4_, 0.5 g/L NH_4_Cl. Additionally, the media contained (per L): 500 µL of 1 g/L sodium resazurin solution, 10 mL of DSMZ 141 trace element solution, 23.81 mL of 1 M sodium bicarbonate, 18.87 mL 1 M sodium carbonate, 5 mL of 40 g/L 2-mercaptoethane sulfonic acid sodium salt (Coenzyme M), and a small spatula tip (1-2 mg) of sodium dithionite as reductant. The pH was adjusted to ∼7.5 and cultures were grown in 9.5 mL of medium in 70 mL serum bottles. Vancomycin and streptomycin (10 µL of 50 mg/mL), 200 µL of 1 M TMA, and 300 µL of carbon source mix [22] were added to each bottle. Bottles were inoculated with 100 µL from the previous culture and grown at 30 °C for approximately one month prior to transfer.

### Methane measurements

Methane was measured via GC-FID as previously described [35] with the following modifications: (i) 1 mL of sample was injected into autosampler vials instead of 250 µL, (ii) a 3 mL plastic BD syringe equipped with a sample lock was used in place of a Hamilton syringe, and (iii) an Rtx-1 GC Capillary column was used instead of a GS-CarbonPLOT column.

### Gene amplicon analysis

16S rRNA gene analysis was performed as described in Krukenberg et al. [36] using Earth Microbiome Project primers 515F and 806R [37, 38]. The mlas-mod-F and *mcrA*-rev-R primers and the touchdown amplification program from Angel et al. 2012 were used to amplify the alpha-subunit of methyl-coenzyme M reductase (*mcrA*) [39]. Further details can be found in the supplementary methods section.

### Metagenomics

A combination of Illumina short-read and Nanopore long read sequencing, following an established workflow [35] with some modifications, was used to obtain the circular genome of *Ca.* M. hillemani str. BB. Fragmented but complete (>99.5 %) genomes were also generated for *Halanaerobium* and *Desulfovermiculus* spp.. The genomes can be found with the following accession numbers for *Ca.* M. hillemani closed genome, *Desulfovermiculus* sp., and *Halanaerobium* sp., respectively: SAMN54119603, SAMN54119604, SAMN54119605. Details on sequencing, assembly, annotation, and genome comparisons are described in the supplementary methods section.

### Salt stress experiment

For methane production and metaproteomics experiments, *Methanohalophilus* enrichment cultures were grown (i) as the control (176 g/L total salts; 50/45/5 of H_2_/N_2_/CO_2_ headspace), (ii) with a 100% N_2_ headspace, (iii) in low salt media (114 g/L salts; 50/45/5 of H_2_/N_2_/CO_2_ headspace), (iv) high salt media (227 g/L salts; 50/45/5 of H_2_/N_2_/CO_2_ headspace), and (v) micro-oxic conditions (176 g/L total salts; 50/45/5 of H_2_/N_2_/CO_2_ headspace plus 1 mL of filter-sterilized room air injected into the headspace every four days, achieving ca. 0.35% O_2_ upon first injection). The ratio of the different salts for all conditions was the same as described for the media above, with each salt being decreased or increased proportionally to achieve the final concentrations. Cultures grown under control conditions were transferred (1:100) into bottles with these conditions. To acclimate the test cultures to stress conditions, they were grown under these test conditions for three transfers before methane data was recorded during the fourth generation. That culture was then used to inoculate a fifth generation, which was harvested in mid-log phase (based on methane production) for the metaproteomics experiment and the 16S rRNA gene community analysis (n=3 replicates for each condition).

### Metaproteomics sample preparation and data analysis

The harvested cultures were sent to the Pacific Northwest National Laboratory (Richland, WA, USA) where 200 µg of protein was used for digestion. MS analysis of the peptides was performed using an Orbitrap Exploris 480 mass spectrometer (Thermo Scientific, Waltham, MA, USA). Peptide matching from the acquired datasets was performed using MSGF+ software [40] against target meta-proteome composed from sequenced genomes (see Supplementary Methods). Peptide abundances, integrated as area under the peptide elution peaks, were produced from raw spectra using in-house developed software MASIC [41]. MASIC peptide abundances were log2 transformed to remove skewness in the distribution of measured abundances. Peptide abundances were corrected for the relative abundance of organisms using the summed area under the peptide curves mapping to each organism. Transformed abundance values of peptides, and later proteins, were normalized using mean central tendency method implemented in InfernoR software [42]. Unless otherwise indicated, all proteomic data presented in this study represents normalized protein abundances. Raw proteomics data has been deposited at massIVE under accession number: MSV000100495. Further details can be found in the supplementary methods section.

### Methane flux measurements

Methane flux data was collected in transects, perpendicular to the shoreline, at Bridger Bay in the south arm of the GSL using A Li-COR 7810 trace gas analyzer (Li-COR, Lincoln, NE, USA). Terrestrial flux measurements were made as previously described [43], but with a one-minute measurement period and a 10 second deadband time. To generate depth-resolved 16S rRNA gene amplicon data, soil cores were collected from within the boundary of the terrestrial collar which registered the highest methane flux. Aquatic flux measurements were done using a custom-made floating flux chamber to take duplicate, one minute, flux measurements. Aquatic fluxes were calculated from the linear change in methane concentration over time, as previously described [44].

## Results and discussion

### Enrichment of a novel methanogen

Microbialite and water samples from Bridger Bay in the South arm of GSL were collected in July 2022, when the water level was at its lowest in recorded history (Supplementary Figure 3). Methanogens were enriched using the growth medium used to isolate *Methanohalophilus mahii* [22]. We increased the salinity of the medium to match GSL water from April 13^th^ 2023, provided a headspace consisting of 50% H_2_, 45% N_2_, and 5% CO_2_, and used trimethylamine (TMA) as the methanogenic substrate. The culture also produced methane from MMA, DMA and methanol, but not from glycine betaine or hydrogen and carbon dioxide (Supplementary Figure 4).

After 21 months of enrichment, 16S rRNA gene amplicon sequencing indicated that the culture consisted of 63% *Methanohalophilus*, 20% *Desulfovermiculus* (a sulfate reducing bacterium - SRB), and 16% *Halanaerobium* (an anaerobic fermenter). An additional 25 taxa contributed to the remaining <1% of the 16S rRNA gene reads. Metagenomic sequencing of this sample with both Illumina and Oxford Nanopore enabled assembly of a circular *Methanohalophilus* genome (2.09 Mb). Sequence coverage indicated that the culture consisted of 80% *Methanohalophilus*, 7% *Desulfovermiculus* sp., 4% *Halanaerobium* sp., with 9% of sequences remaining unbinned. In addition to these three assembled genomes, the metagenome contained 16S rRNA gene sequences belonging to *Vibrio cholerae*, a *Desulfohalobium* sp., and an *Izimaplasma* sp.. Attempts to isolate the methanogen using dilution to extinction, colony-picking from solid media, and additional antibiotics were not successful.

### Characterization of the novel methanogen as *Ca.* Methanohalophilus hillemani

This newly assembled *Methanohalophilus* genome shares less than 95% ANI with all other genomes within its genus, indicating a novel species [45] and only 91.2% with *M. mahii* (Figure 1A). The 16S rRNA gene is less than 99.3% identical to other species, noticeably lower than other *Methanohalophilus* species to each other (Supplementary Figure 5). We propose the name *Candidatus* Methanohalophilus hillemani strain BB for this archaeon (see etymology section in the supplementary online materials).

**Figure 1:**
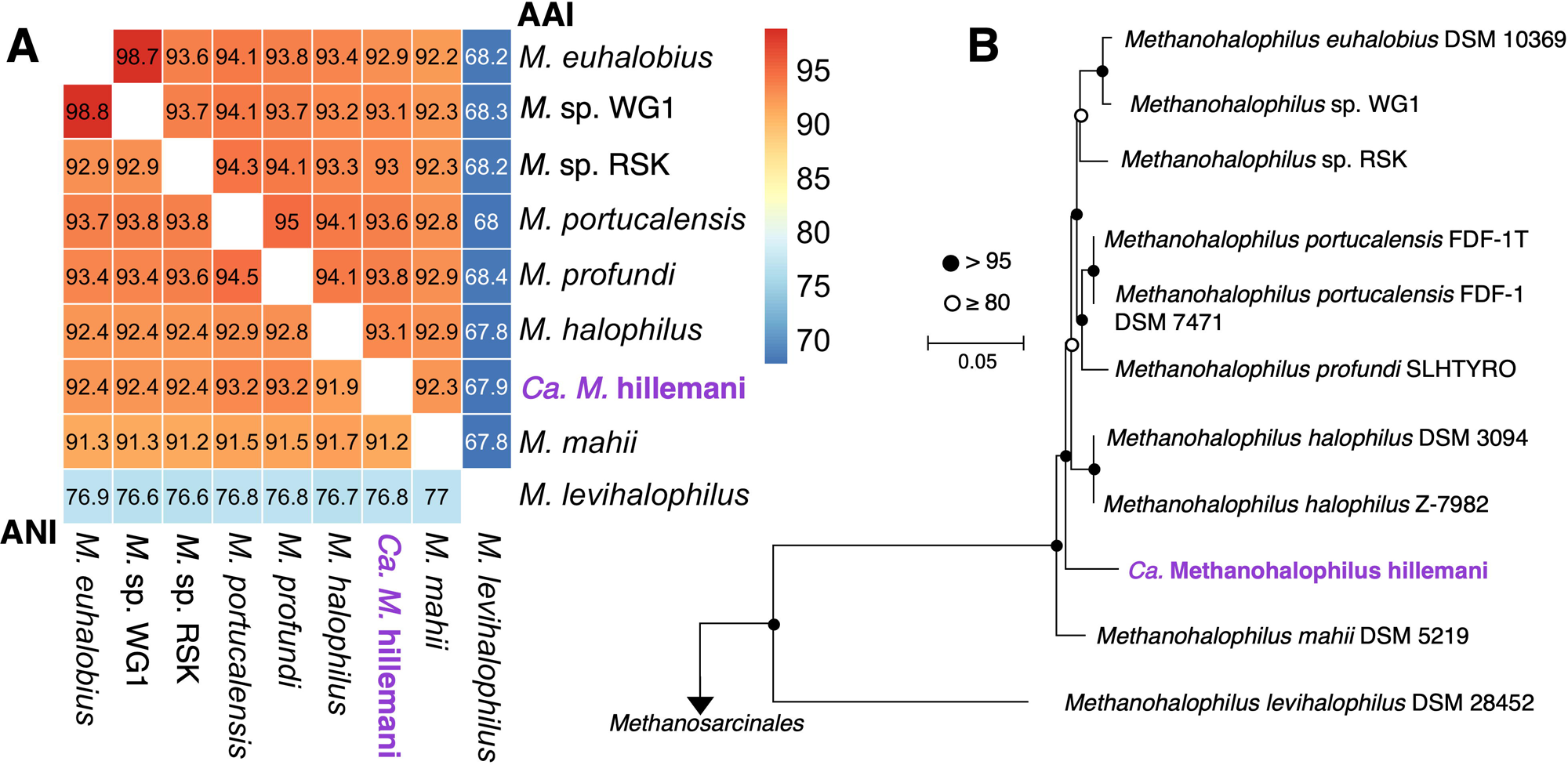
*Ca.* Methanohalophilus hillemani represents a novel species within the genus *Methanohalophilus*. (A) Pairwise average nucleotide identity (ANI) and average amino acid identity (AAI) on bottom left and top right, respectively, demonstrating *Ca.* M. hillemani represents a new species within the *Methanohalophilus* genus. (B) Maximum likelihood tree of 65 aligned and concatenated single-copy *Methanohalophilus* proteins constructed with FastTree v2.1.11 [46] and 1,000 bootstraps. Circles represent bootstrap values. Nodes with <50 support were collapsed. Accession numbers for the other *Methanohalophilus* species can be found in SI Table 1.

Comparing protein-coding sequences encoded by *Ca.* M. hillemani to all other *Methanohalophilus* spp. showed it shares the major characteristics of the genus, including methyl-dismutating methanogenesis from MMA, DMA, TMA, and methanol [23, 28, 34 (SI Table 2)]. Like the other species, *Ca.* M. hillemani lacks hydrogenases that would be required for hydrogenotrophic methyl-reducing methanogenesis. Other conserved functions across the genus include glycine betaine transport and synthesis, synthesis and usage of the 22^nd^ amino acid, pyrrolysine (Pyl), and O_2_ tolerance via thioredoxins, peroxidases, superoxide reductases, and peroxiredoxin. Unique genetic features of *Ca.* M. hillemani as compared to other members of the genus *Methanohalophilus* are discussed in the Supplementary Results section.

### Increasing salinity in the Great Salt Lake

The biomass that led to culturing *Ca.* M. hillemani was collected from the GSL in July 2022, when the lake was at its lowest recorded level (Supplementary Figure 3). Additionally the sample came from Bridger Bay, which was cut off from the main lake by low water levels, leading to a salinity much higher (∼24%) than the rest of the south arm of the GSL (∼17%). Na^+^ and Cl^-^ concentrations at this time were 3.6 and 4.2 M, respectively, more than twice the concentrations of these ions measured by Post et al. in 1977 (1.6 and 1.8 M) [47]. In contrast, *Methanohalophilus mahii* was isolated in 1985 [22], right before GSL reached recorded high water levels (Supplementary Figure 5). This led us to the hypothesis that *Ca.* M. hillemani may be adapted to higher osmotic stress levels as compared to *M. mahii*.

### Methane production under salt stress

To understand how stressors in the GSL ecosystem might impact *Ca.* M. hillemani, control cultures were compared to those incubated under low salt, high salt, 100% N_2_, and micro-oxic conditions (see methods, above). Proportions of major salt ions in all conditions matched the GSL, which contrasts previous work on the genus that had only modulated NaCl concentration [22, 29, 48]. The lag time for methane production increased slightly with increasing salinity (Figure 2A). Nevertheless, the overall methane yield for *Ca.* M. hillemani cultures was consistent across all conditions tested (Figure 2A). Mass balance of the methyl groups provided via TMA shows that after 16 days 68 - 77% was converted to methane, consistent with the 75% value for methyl-dismutating methanogenesis that has been reported for some euryarchaeotal methanogens [49, 50].

**Figure 2:**
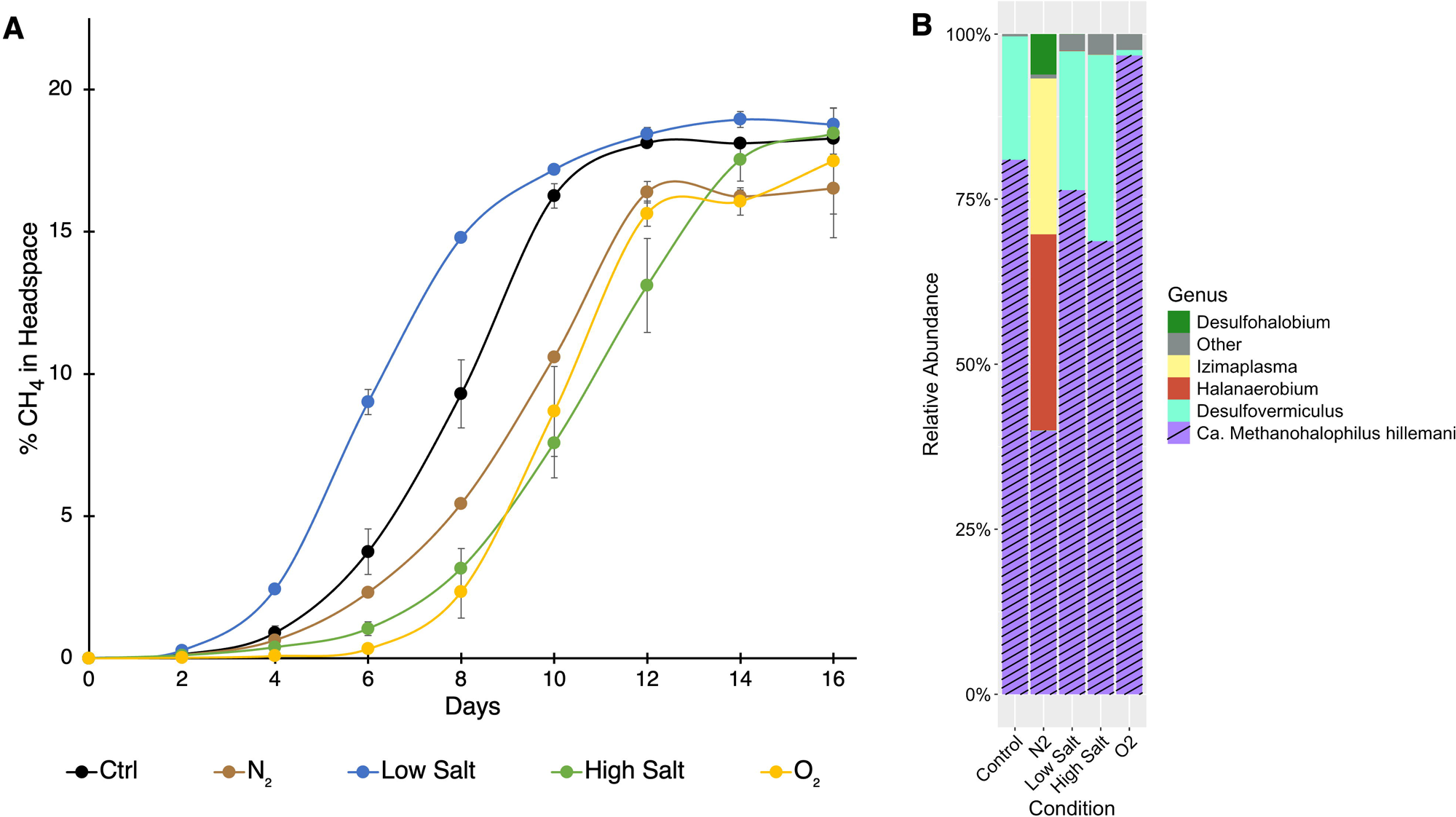
*Ca.* Methanohalophilus hillemani produces similar amounts of methane under a variety of conditions. (A) Methane production of the *Methanohalophilus* enrichment culture grown under various stress conditions. Error bars (where visible) represent the standard deviation of triplicate cultures. (B) Community composition of same samples that were processed for metaproteomics, as assessed by 16S rRNA gene amplicon sequencing. Amplicon sequence variants that constituted less than 0.5% of the total community were clustered into the “Other” category.

16S rRNA gene amplicon data showed that *Ca.* M. hillemani was the most abundant member in all conditions (Figure 2B); however, the community composition of these cultures differed significantly from the 3-species enrichment indicated by the metagenome. The control, low salt, and high salt conditions contained primarily (>68%) *Ca.* M. hillemani, while D*esulfovermiculus* was the only other organism with a relative abundance over 0.5%. Over 96% of amplicons in the O_2_ condition belonged to *Ca.* M. hillemani. By contrast, in the N_2_ condition 40% of amplicons belonged to *Ca.* M. hillemani and less than 0.1% to *Desulfovermiculus*. An *Izimaplasma* species that was absent from all other samples represented 24% of the community, and the *Halanaerobium* species that made up <0.1% of amplicons in all other conditions, constituted 30% in the N_2_ condition. Despite differences in culturing conditions and community composition, especially in the N_2_ cultures, all conditions yielded the same amount of methane.

### Metaproteomics

To understand the biological basis for *Ca.* M. hillemani’s consistent methane production across stressful conditions, we performed metaproteomics. Triplicate cultures were harvested in mid-log phase and peptides from cultures of each condition were mapped to predicted proteins from the assembled genomes. Because no bins could be retrieved from *Izimaplasma* and *Desulfohalobium* from the N_2_ condition, we were not able to map peptides to these organisms.

### Stress conditions do not impact methanogenesis protein expression

For all conditions, the proteins involved in the methyl-dismutating methanogenesis pathway were among the top 10% of *Methanohalophilus* proteins expressed (Supplementary Figure 6). The proteins responsible for the transfer of a methyl group to coenzyme M and its subsequent reduction to methane were all among the top 1%, with one exception (methylcobamide:CoM methyltransferase – in the top 1.1% of the N_2_ condition). The only methanogenesis-related protein that was differentially expressed was an MMA-methyltransferase protein found in the control condition but not in the O_2_ condition (Figure 3). However, *Ca.* M. hillemani encodes two copies of this protein, and the other copy was in the top 1% of proteins expressed in the O_2_ condition, suggesting that the methanogenic pathway was not impacted.

**Figure 3:**
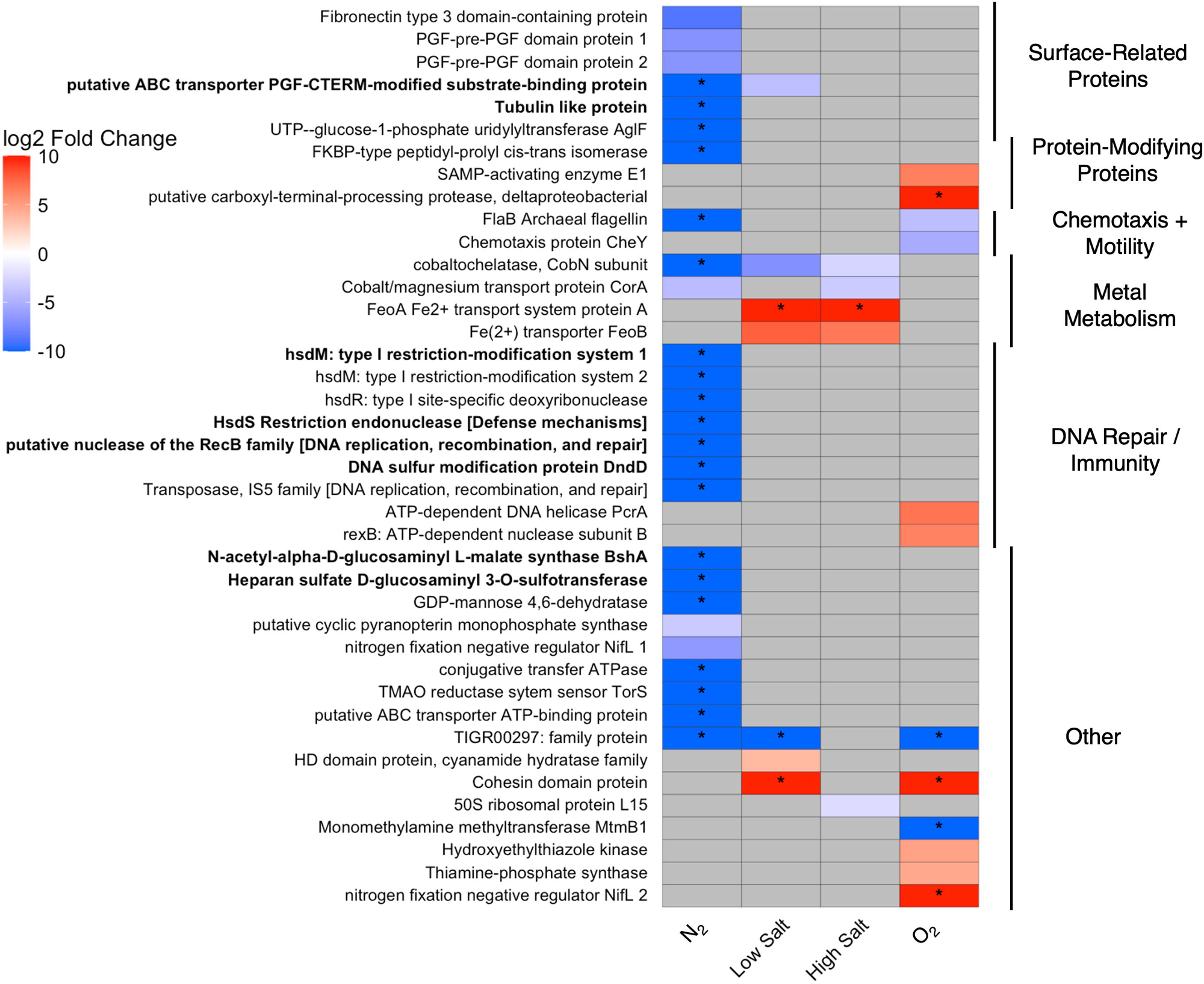
*Ca.* Methanohalophilus hillemani differential protein expression between control and test conditions. Maximum differentially expressed proteins occurred between the control and N_2_ conditions, all of which were more highly expressed in the control. Down-regulated proteins in the N_2_ condition included proteins involved in immune response, metal metabolism, and the cell-surface. Proteins shown in this figure meet three criteria: (i) above median normalized protein expression for that condition; (ii) statistically significantly differentially expressed (p < 0.05); and (iii) function could be predicted (*i.e.* hypotheticals not included). A negative log fold change indicates the protein was more abundant in the control, positive indicates more abundant in the test condition. Asterisks in the boxes indicate that the protein was not detected in either that test condition or the control and a log2 fold change of 10 or -10 was imputed. Bolded proteins indicate that the coding sequence for this protein was unique to *Ca.* M. hillemani and no homologs were identified in other *Methanohalophilus* genomes. Proteins with grey boxes do not indicate absence in a specific condition, but that the protein did not meet all three criteria listed above. See Supplementary Tables 3-7 for raw values.

It was previously reported that in response to long-term high and low-salt stress, *Methanohalophilus portucalensis* transcripts for methylamine methyltransferases and corrinoid proteins were upregulated [48]. Conversely, we show that the *Ca.* M. hillemani methanogenic proteome is highly and uniformly expressed, regardless of high or low salt concentrations. One possible explanation for the lack of observed methanogenic protein expression response, is the high basal expression of energy-conservation proteins (Supplementary Figure 6). This may be advantageous in an environment with constant stressors that require large amounts of energy to tolerate (*i.e.* osmotic and oxidative stress). This strategy could be especially feasible in the GSL, where methylated amines are likely always in high abundance due to the production of a metabolic precursor, glycine betaine [51], as a compatible solute.

Another possibility is that methanogenic protein activity, as opposed to abundance, is post-translationally regulated in response to stress. An investigation of protein phosphorylation in *Methanohalophilus portucalensis* found that 26% of the phosphorylated proteins were involved in methanogenesis, energy conservation, and osmolyte biosynthesis [52]. That study found that a mutated glycine sarcosine N-methyltransferase (responsible for glycine betaine synthesis), modified to act like a phosphorylated version, had reduced activity that resulted in lower salinity tolerance when expressed in *E. coli*. While they did not have analogous data for phosphorylated methanogenic proteins, they demonstrate that *Methanohalophilus* proteins can be regulated beyond the level of protein expression. The transcriptomic and proteomic findings discussed here, together with the lack of differential expression of methanogenic proteins, suggests that future work on the *Methanohalophilus* genus should examine forms of regulation downstream of protein expression.

### Limited number of proteins differentially expressed in response to changing salinity

Contrary to our expectations, low and high salt conditions resulted in relatively few proteins that were both highly and differentially expressed, compared to the control (Figure 3). Both high and low salt conditions showed two upregulated proteins from an iron transport system, FeoA and FeoB, of which FeoA was not detected in the control. Of the many DNA repair proteins encoded by *Ca.* M. hillemani, none were both upregulated and expressed above median protein expression in the high salt condition. The lack of DNA-repair protein expression in response to high salinity was unexpected [53], though basal expression of several of these proteins was observed in the control. Alternatively, the need for DNA repair may be mitigated by osmo-tolerance via high expression of compatible solute. pathways. Indeed, glycine betaine synthesis and import genes are in the top 10% of expressed proteins in the low-salt, high-salt, and control conditions. As discussed above, post-translational regulation could also play a role in osmo-tolerance without requiring differential expression of proteins [52].

Protein expression was more affected by the O_2_ growth condition. Two proteins associated with protein modification and two DNA-repair proteins were up-regulated and expressed above the median (Figure 3). An additional three DNA-repair proteins were up regulated but not expressed above the median. Thiamine is known to confer antioxidative stress properties; two proteins involved in thiamine biosynthesis, hydroxyethylthiazole kinase and thiamine phosphate synthase, were upregulated in the O_2_ condition. Surprisingly, none of the oxygen detoxification genes were upregulated. Constitutive expression could also explain this lack of response to oxygen—superoxide reductase, peroxiredoxin, and catalase-peroxidase are all in the top 10% of expressed proteins in the control.

### Unique *Ca.* M. hillemani proteins are differentially expressed in the N_2_ condition

When comparing the number of differentially expressed proteins between the control and test conditions, the N_2_ condition resulted in twice as many differentially expressed proteins (Supplementary Table 8). This was consistent when filtering for only the proteins that were expressed above the median for each condition or control (Supplementary Table 9). Notably, all 26 differentially and highly expressed *Ca.* M. hillemani proteins in the N_2_ condition were less abundant than in the control (Figure 3). Additionally, eight of these 26 differentially and highly expressed proteins are unique proteins to *Ca.* M. hillemani, among the *Methanohalophilus* genus (see Supplementary Results). However, none of these unique and highly expressed proteins were detected in the N condition; in contrast, all eight were found in every other condition, with seven expressed above the median. We hypothesize that these distinct protein expression patterns in the N_2_ condition are due to the differences in community composition (Figure 2B).

### Metal uptake, DNA repair, and immunity down-regulated in the N_2_ condition

Down-regulated functions in the N_2_ condition included immune response and DNA repair, cell-surface features, and metal metabolism (Figure 3). A cobaltochelatase and a cobalt/magnesium transporter were not detected and down-regulated 19-fold, respectively, in the N_2_ condition (Figure 3). Additionally, one iron transporter was down regulated 29-fold and two other iron transporters, detected in all other conditions, were absent from the N_2_ condition.

Highly expressed immune-response and DNA-repair proteins were noticeably absent from the N_2_ condition. Seven proteins in this category (Figure 3) were undetectable in the N_2_ condition yet were expressed above the median in all other conditions. Eleven additional proteins involved in immune response or DNA repair were also detected in every condition except for N_2_. These included two CRISPR-related proteins, two proteins involved in the sulfur-modification of DNA [54], four DNA helicases, two DNA repair proteins, and an additional copy of the type 1 restriction-modification system protein, HsdS.

### *Ca.* M. hillemani is adapted to sulfide stress and competition with *Desulfovermiculus*

The genomic and proteomic data do not support the hypothesis that osmotic tolerance distinguishes *Ca.* M. hillemani and *M. mahii*. Both species are genetically capable of synthesizing and taking up the compatible solute, glycine betaine, and both encode ion transporters to maintain osmotic balance. While *Ca.* M. hillemani encodes unique proteins involved in DNA repair and immune response, it encodes no unique proteins predicted to specifically promote osmotolerance (Supplementary Table 10). In fact, differing salt concentrations resulted in a relatively small proteomic change in this halophilic methanogen.

Instead, the main proteomic differences were observed from the N_2_ condition. Immune-response and DNA-repair proteins, including proteins unique to *Ca.* M. hillemani, were highly expressed in all conditions except for the N_2_ condition. It was previously shown that DNA-repair transcripts were upregulated by methanogens in response to the presence of sulfide [55]. Additionally, high levels of sulfide can be detrimental to methanogens [56, 57] and known to damage DNA [58]. Sulfide is produced by SRBs using H_2_ as an electron donor, and notably, all conditions in our study, except for the N_2_ condition, contained a headspace with 50% H_2_. While we did not take sulfide measurements during the proteomics experiment, conditions with H_2_ in the headspace yielded cultures with a 15-500 times higher relative abundance of the SRB *Desulfovermiculus*, compared to the N_2_ condition (Figure 2B). A different SRB, *Desulfohalobium,* was present in the N_2_ condition at lower relative abundances. However, since no bin for this genus was retrieved from our metagenome, we were unable to map proteins to this organism. Together, these results indicate that either high levels of sulfide or the presence of the specific SRB *Desulfovermiculus* induced a distinct proteomic response by *Ca.* M. hillemani. Unique immune-response and DNA-repair proteins expressed by *Ca.* M. hillemani in the presence of *Desulfovermiculus* potentially represent novel adaptations to protect or repair DNA from SRB-related damage.

Additionally, reduced expression of metal uptake and chelation by *Ca.* M. hillemani in the N_2_ condition, compared to all other conditions, suggests competition for metals with *Desulfovermiculus*. Cobalt is crucial for methyl-dismutating methanogenesis as it is the cofactor in corrinoid proteins used to transfer methyl groups to MCR (Supplementary Figure 6) [59]. However, the conversion of sulfate to sulfide by SRB can directly diminish the availability of cobalt through Co^2+^ - sulfide co-precipitation [60]. Additionally, SRB utilize cobalt as an enzymatic cofactor for various metabolisms, including dissimilatory sulfate reduction [61, 62], which could lead to competition. Indeed, *Desulfovermiculus* expresses two cobaltochelatases in the control condition, one just below median normalized expression and the other well above. Tolerating sulfide stress and competing against *Desulfovermiculus* for cobalt may be important factors impacting *Ca.* M. hillemani physiology when H_2_ is available.

### *Ca.* M. hillemani uses pyrrolysine sparingly

Methylated amine-utilizing methanogens are known to encode the 22^nd^ amino acid, pyrrolysine (Pyl), which plays an important role in methylamine activation in tri, di, and monomethylamine methyltransferases (MttB, MtbB, MtmB, respectively) [63–65]. Indeed, *Ca.* M. hillemani encodes the genes required for Pyl synthesis (PylBC) and uses (Pyl-tRNA ligase). An mRNA secondary structure element, PYLIS (analogous to SECIS for selenocysteine), has been suggested to enable Pyl insertion into proteins [66, 67] and PYLIS elements were annotated within *Ca.* M. hillemani’s methylated-amine methyltransferase genes. Accordingly, Pyl residues were identified in peptidesmapping to all three types of methylated-amine methyltransferases encoded by *Ca.* M. hillemani.

Kivenson *et al*. have recently proposed that some methanogens, including *Methanohalophilus mahii*, *M. euhalobius*, and *M. levihalophilus,* have reprogrammed the TAG codon to universally incorporate Pyl, instead of being read as a Stop-codon, across the proteome [68]. While the authors confirmed their hypothesis for a marine and a human-gut-associated methanogen, experimental data on halophilic methanogens was not yet available. Moreover, how and if microbes regulate the incorporation of Pyl at TAG codons is a topic of current debate [68, 69]. Having already identified Pyl residues in peptides mapping to methyltransferase proteins, we used Kivenson’s modification to Prodigal [68], as well as our own modifications to Prokka, to re-predict and annotate genes for *Ca.* M. hillemani. This resulted in 95 predicted proteins containing an in-frame TAG codon. In this reanalysis, we found peptide hits from predicted-Pyl-containing proteins downstream of the predicted-Pyl residue. This could suggest that the TAG codon is resulting in a Pyl residue, not being read as a Stop-codon. However, for non-methyltransferase proteins predicted to contain pyrrolysine, only 7 of the 3,463 (0.2%) detected peptides associated with those proteins contained Pyl. In contrast, 2,920 out of the 27,114 (10.7%) peptides associate with methyltransferase proteins contained Pyl. The low percentage of non-methyltransferase peptides identified as containing Pyl could indicate that TAG codons, in contrast to genomic predictions for the genus [68], are primarily read as Stop in *Ca.* M. hillemani.

### High methane flux from the Great Salt Lake

To assess methanogen activity *in situ*, we measured methane flux from the GSL. We focused on Bridger Bay, where the *Ca.* M. hillemani inoculum was obtained (Supplementary Figure 2). Methane fluxes were very high but also variable, with fluxes reaching 3 mmol CH_4_/m^2^/day observed across sampling dates (Figure 4A, Supplementary Figure 7). These fluxes are comparable to high-methane flux environments such as thawing permafrost [43, 70] and freshwater wetlands [71]. The highest flux measured was 377 mmol CH_4_/m^2^/day (Supplementary Figure 8), which greatly exceeds values reported from either permafrost or wetlands.

**Figure 4:**
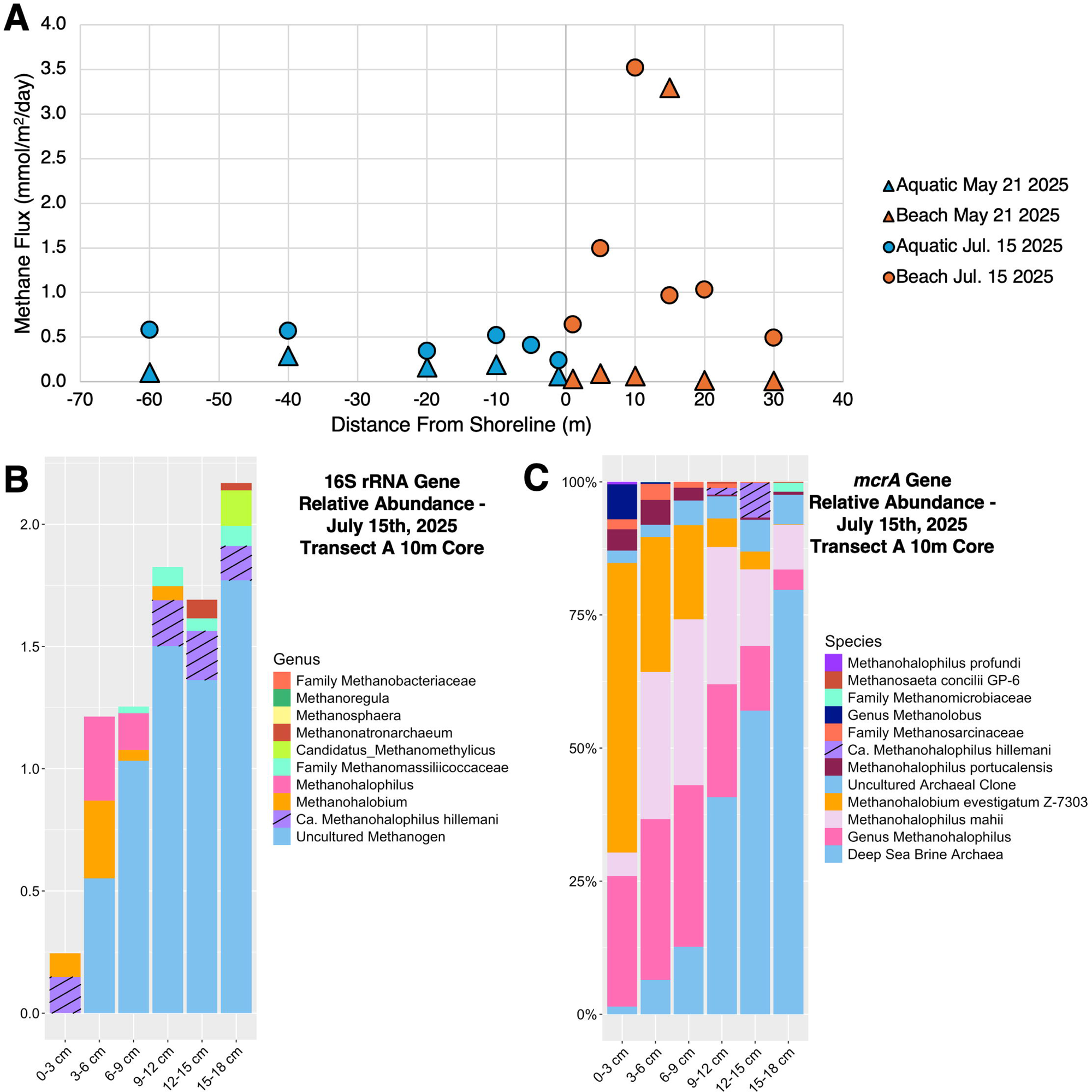
Methane flux and methanogenic community from Bridger Bay on the Great Salt Lake. (A) Methane fluxes measured from soil within Transect A at Bridger Bay in May and July of 2025 showing a zone of high methane flux 10-15 m from the shoreline. (B) Relative abundance of different methanogens in a sediment core taken from the Transect A 10 m flux site from July 15, 2025 based on 16S rRNA gene amplicon sequencing. For most taxa, taxonomic classification was done at the genus level. *Ca.* M. hillemani was manually categorized based on a one-nucleotide difference from other *Methanohalophilus* 16S rRNA genes in their amplified region. Sequences that could not be assigned a genus are labeled “Family” or “Order” prior to the name. “Uncultured Methanogens” (possibly members of the order *Methanofastidiosales*, see main text) were the most abundant sequences. Only methanogen sequences are shown. (C) *mcrA* gene amplicon relative abundance in the same core samples as (B), corroborating the presence of *Ca.* M. hillemani. Other *Methanohalophilus mcrA* gene sequences were also present, including some that could not be classified to the species level. For most taxa, taxonomic classification was done at the species level. For taxa that could not be assigned to species, “Family” or “Genus” is included in the name.

Fluxes were measured in transects perpendicular to the water’s edge, including both terrestrial and aquatic measurements. We repeatedly observed a zone of elevated terrestrial methane flux between 5 and 15 meters from the shoreline (Figure 4A). This was counterintuitive, as we expected that the most anoxic and methanogen-conducive conditions to be right at the shoreline. However, this trend was observed across sampling dates and transects (Supplementary Figure 7).

Our results contrast the only other study that measured methane flux from the GSL [20], which observed similar aquatic fluxes (0.42 +/- 0.08 mmol CH_4_/m^2^/day), but much lower terrestrial fluxes (0.47 +/- 0.1 mmol CH_4_/m^2^/day). Their findings of similar terrestrial and aquatic fluxes and minimal flux variability between sites did not corroborate our results. We hypothesize that this disparity is due to the difference in spatial scales for our two studies. Cobo *et al*. [20] examined differences between basins, over kilometer scales. Their transects included steep vertical gradients over a kilometer from the contemporary GSL shoreline. Our measurements were on the meter scale and centered on the contemporary lake shore. It is only with our higher spatial resolution (and smaller scope), that we observed extremely high methane fluxes and flux variability. Together, Cobo’s and our results show that the GSL does have extremely high methane flux across seasons; however, the zone of highest flux is located just meters from the current shoreline.

### Novel methanogens likely contribute to Great Salt Lake methane flux

To examine the ecological relevance of the novel methanogen in the GSL ecosystem, we performed 16S rRNA and *mcrA* gene amplicon sequencing on samples from a soil core taken from the transect A 10-meter sample, where extremely high and variable methane flux was observed (Supplementary Figure 8). The 16S rRNA gene amplicon data revealed that across all depths, *Ca.* M. hillemani was the second most abundant methanogen, while other *Methanohalophilus* sequences were the fourth most abundant (Figure 4C). *Desulfovermiculus* was the most abundant genus of SRB (Supplementary Figure 9). *Ca.* M. hillemani was similarly abundant across all depths, and it represented the largest fraction of the methanogenic community near the surface. This suggests some level of O_2_ tolerance, consistent with the higher relative abundance of *Ca.* M. hillemani in the O_2_ condition of our culture experiments (Figure 2B). The most abundant methanogens, by far, were represented by a group of 11 poorly characterized sequences. A classifier trained on the ARB-silva 16S rRNA gene database could only identify these sequences as members of the order *Methanofastidiosales* [72], while BLAST only retrieved “uncultured archaeon clone” sequences. The relative abundance of all methanogens increased with depth, ranging from ∼0.25% at the surface to over 2% of the total community at the 15-18 cm depth.

The most abundant *mcrA* gene amplicon sequences were also poorly characterized – their top BLAST hits were to sequences recovered from deep sea anoxic brines (Figure 4C). *Methanohalophilus* sequences that could not be assigned to a specific species were the next most abundant. Of the *Methanohalophilus* sequences that were identified to the species level, *M. mahii* was the most abundant, followed by *M. portucalensis* and *Ca.* M. hillemani. Like the 16S rRNA gene amplicon data, the poorly characterized sequences increased in relative abundance with increasing depth. In both the 16S rRNA and *mcrA* gene amplicon datasets, the most abundant methanogenic sequences were poorly characterized, indicating that novel methanogens may be the dominant community in the high-methane-flux GSL soil.

### Conclusion

*Ca.* Methanohalophilus hillemani is a novel methanogen, collected from the south arm of the GSL when its salinity was at an all-time high. Metaproteomic analysis of this organism showed that methanogenesis and osmo-tolerance proteins had high basal expression. The expression of these proteins was not impacted by salinity changes, suggesting that future proteomic studies of the *Methanohalophilus* genus should focus on post-translational modification and other forms of regulation, for example on protein activity or metabolome level. Large differences in the expressed proteome were instead observed in the N_2_ condition, which lacked a large population of the SRB, *Desulfovermiculus*. In conditions under which *Ca.* M. hillemani and *Desulfovermiculus* were both present, the methanogen highly expressed metal transporters and chelatases. We hypothesize that, with sufficient H_2_, SRB directly compete for and reduce the availability of trace metals, forcing *Ca.* M. hillemani to invest in uptake of metals vital to its energy metabolism. Coculturing of the two also led to the upregulation of unique immune-response and DNA-repair proteins by *Ca.* M. hillemani, indicating the adaptation of novel strategies to tolerate stress and/or *Desulfovermiculus*. High methane flux was measured from the shore of the GSL, and amplicon data of a soil core identified *Ca.* M. hillemani to be present at significant levels, confirming the ecological relevance of this novel methanogen. In summary, this study cultured a novel methanogen from the GSL, demonstrated that its physiology may not be controlled as much by changes in salinity as by community interactions, and identified it in GSL soils exhibiting high methane flux.

## Acknowledgements

This work was supported by the NASA Exobiology award 80NSSC21K0487 (to T.R.M.) and Department of Energy, Office of Science, Biological and Environmental Research award DE-SC0025661 (to R.H.). A portion of this research was performed under FICUS award 509790 (DOI:10.46936/fics.proj.2023.60847/60008905) and used resources at the Environmental Molecular Sciences Laboratory, a DOE Office of Science User Facility sponsored by the Biological and Environmental Research program under contract no. DE-AC05-76RL01830. Battelle operates PNNL for the DOE under contract DE-AC0576RLO01830. We thank Bonnie Baxter (Westminster University) for her guidance on field sampling, Angel Nieto and Ciara Pike for assistance during field work, Veronika Kivenson (UC Berkeley) for helpful discussions on pyrrolysine and alternative codon use, and the laboratory of Samantha Joye (University of Georgia) for help in quantifying major ions in GSL water. Finally, we thank the family of the late Dr. Maurice Hilleman for allowing us to name the newly cultured methanogen after him in recognition of his life-saving contributions to microbiology, vaccine development, and public health (see Supplementary Etymology).

## Contributions

Conceptualization: W.C.C., T.R.M. and R.H.

Methodology: W.C.C., T.R.M. and R.H. Validation: W.C.C., Z.J.J., N.T., T.R.M., and R.H.

Formal analysis: W.C.C., Z.J.J. and N.T.

Investigation: W.C.C., C.D.N., S.T. and R.L.

Resources: T.R.M. and R.H.

Data curation: W.C.C., Z.J.J., and N.T.

Writing–original draft: W.C.C.

Writing–review and editing: W.C.C., Z.J.J., N.T., T.R.M., and R.H.

Visualization: W.C.C. and Z.J.J.

Supervision: T.R.M. and R.H.

Project administration: T.R.M. and R.H.

Funding acquisition: T.R.M. and R.H.

## Supplementary figures

**Supplementary Figure 1.**
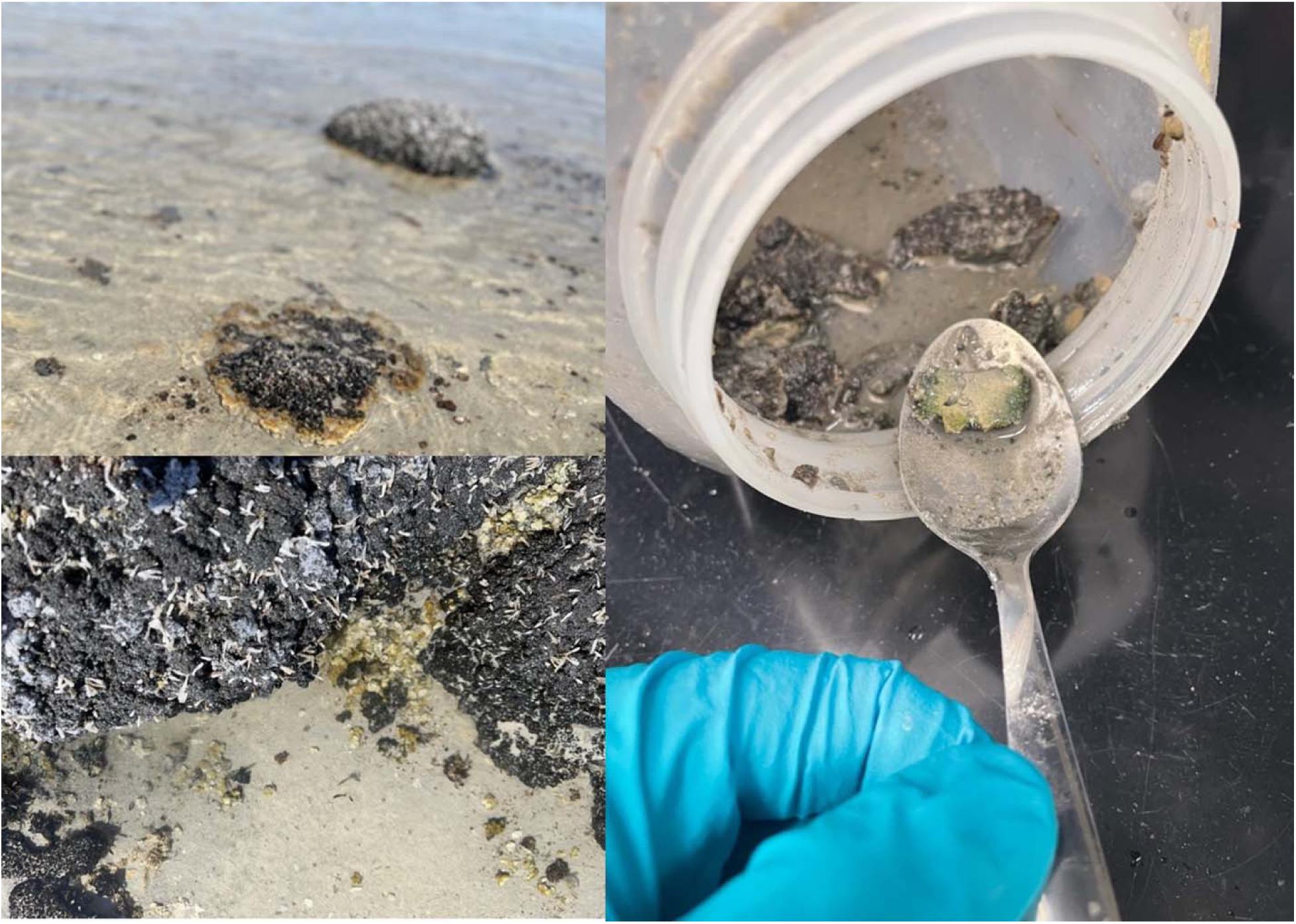
Microbialite material from which the enrichment culture was started. Material shown in the spoon was the exact material from which *Ca.* Methanohalophilus hillemani was enriched.

**Supplementary Figure 2:**
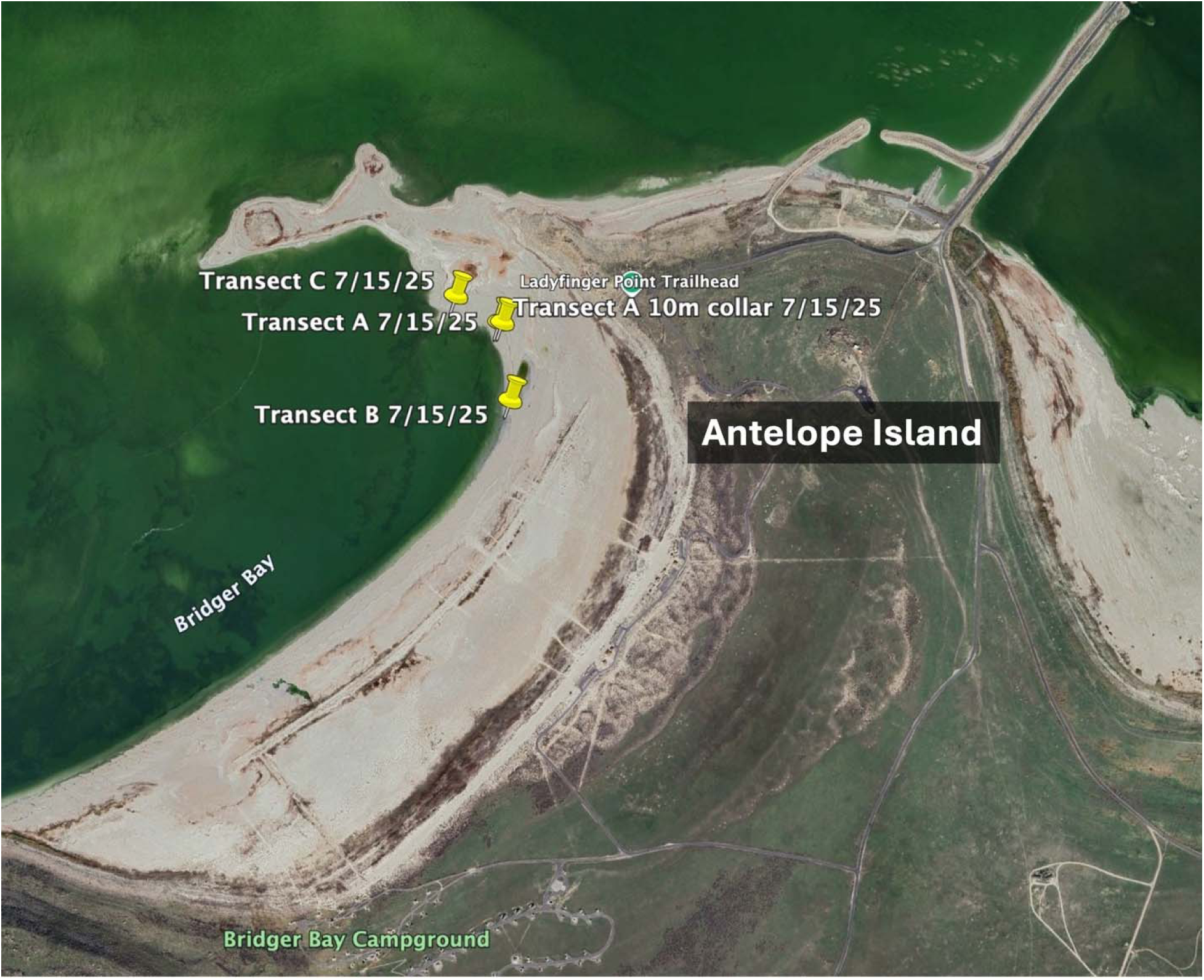
Methane flux transects from July 15^th^ 2025. Locations of the transects A, B and C as sampled on July 15^th^ 2025. The pinned location represents the shoreline (1m) flux measurement. Measurements extended further into the shore and out into the water, perpendicularly, from this point.

**Supplementary Figure 3:**
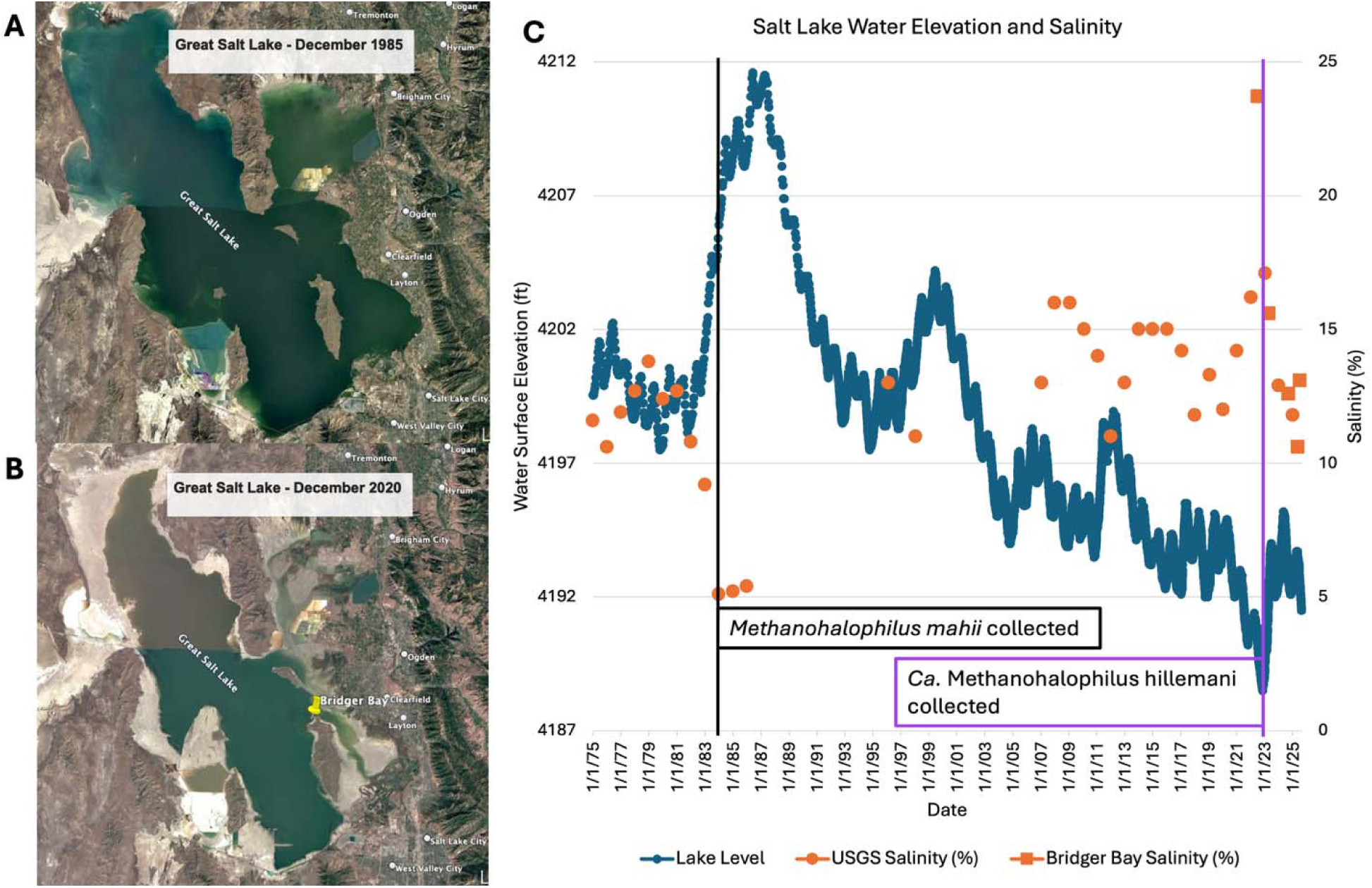
Change in the Great Salt Lake during past half century. Google Earth satellite imagery from the Great Salt Lake near its recorded high lake level in 1986 (A) and near its recorded low in 2022 (B). (C) Lake level and the reciprocal salinity varied substantially at the times when *M. mahii* and *Ca.* M. hillemani were cultured. Lake level data was retrieved for Saltair Boat Harbor from the USGS website. “USGS Salinity” data was only available on the website starting in August of 1995. Salinity data prior to this was retrieved from Arnow and Stephens 1990. Salinity data from Bridger Bay was collected in this study and measured using a refractometer.

**Supplementary Figure 4:**
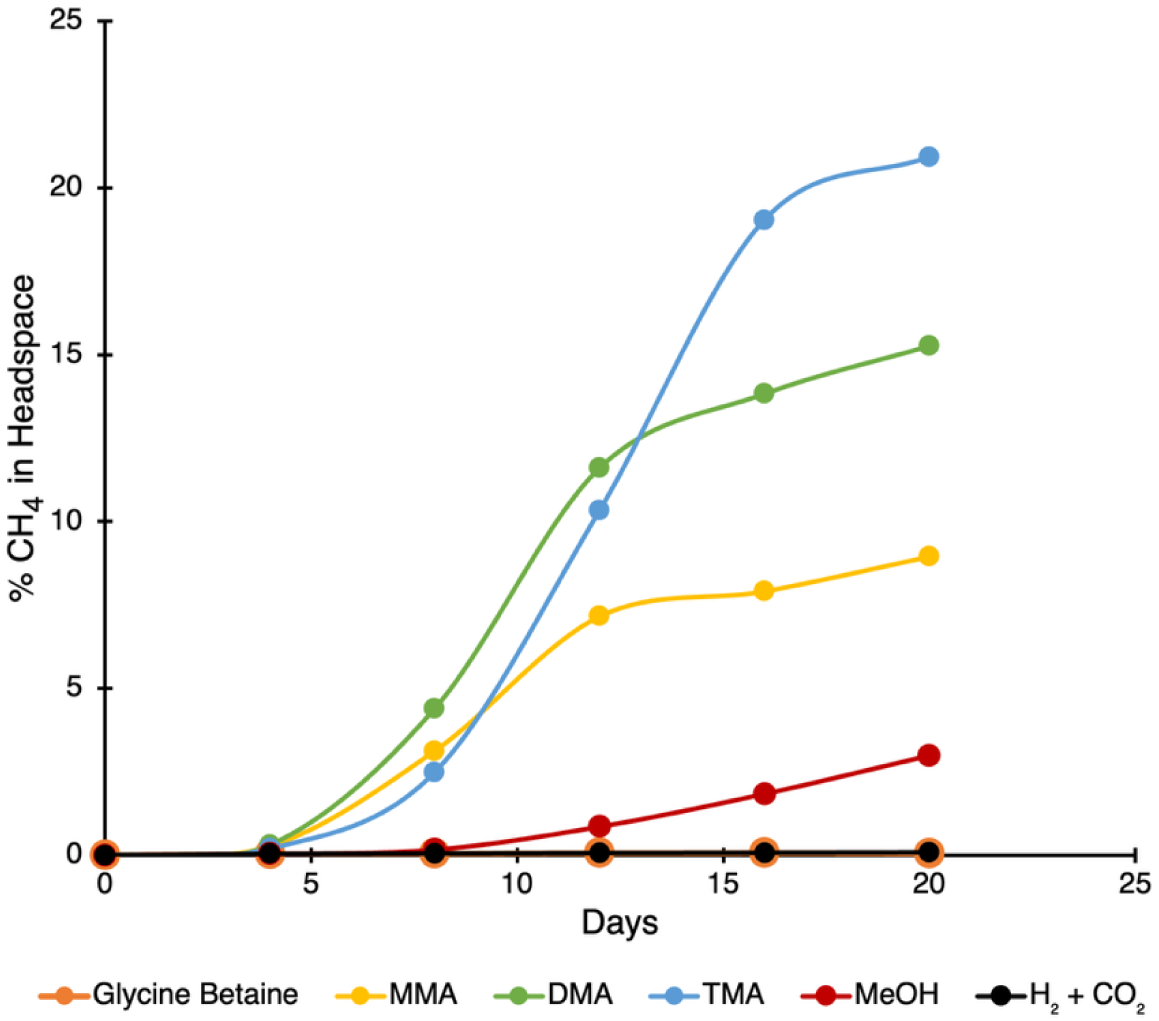
Methane production from various substrates. The GSL methanogenic enrichment culture produced methane from MMA, DMA, TMA, and methanol, but not from glycine betaine, or H_2_ and CO_2_.

**Supplementary Figure 5:**
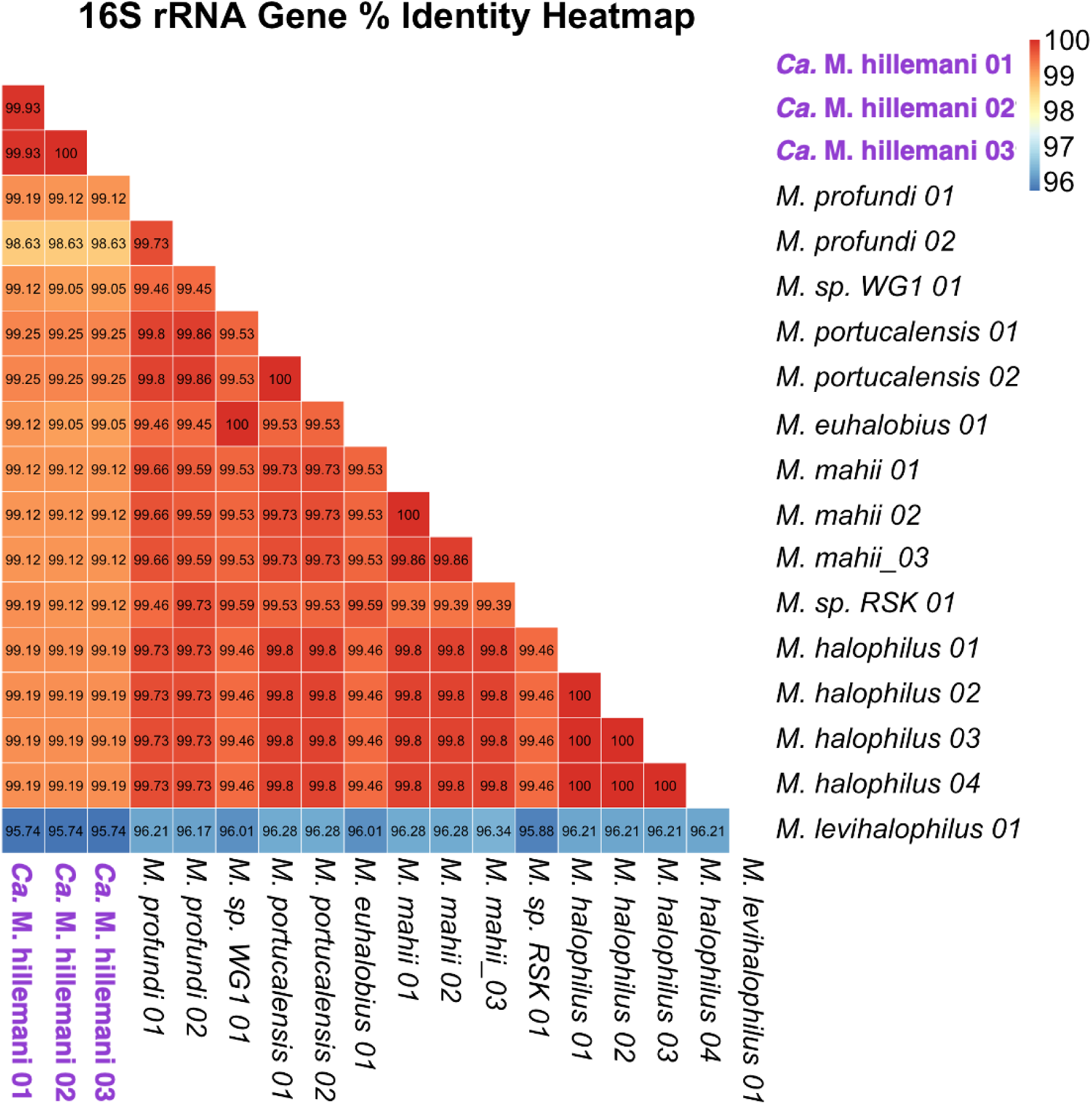
Heatmap of 16S rRNA gene similarity. Heatmap showing the percent identity of the 16S rRNA genes in the *Methanohalophilus* genus. All copies from each genus are shown, should they have multiple copies.

**Supplementary Figure 6:**
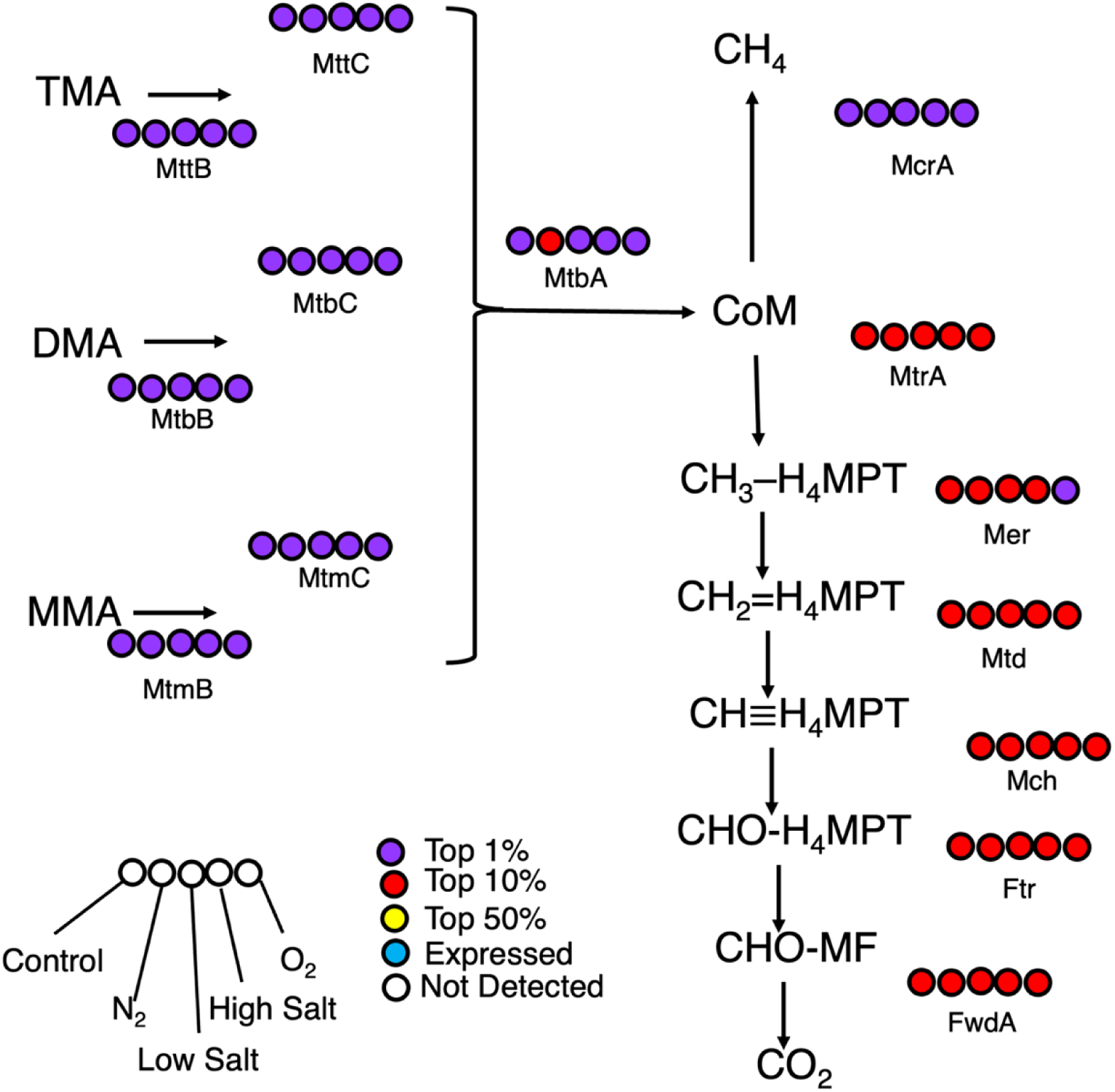
Expression of proteins involved in methyl-dismutating methanogenesis. For each condition the normalized protein values were compared to others within that condition to assess the percentile of expression. In the case of duplicate proteins, the most highly expressed copy was used for this figure. Abbreviations: MttB, Trimethylamine Methyltransferase; MttC, Trimethylamine Corrinoid Protein; MtbB, Dimethylamine Methyltransferase; MtbC, Dimethylamine Corrinoid Protein; MtmB, Monomethylamine Methyltransferase; MtmC, Monomethylamine Corrinoid Protein; MtbA, Methylcobamide:CoM methyltransferase; McrA, Methyl Coenzyme M Reductase Subunit α; MtrA, Tetrahydromethanopterin S-methyltransferase subunit A; Mer, 5,10-methylenetetrahydromethanopterin reductase; Mtd, F420-dependent methylenetetrahydromethanopterin dehydrogenase; Mch, Methenyltetrahydromethanopterin cyclohydrolase; Ftr, Formylmethanofuran--tetrahydromethanopterin formyltransferase; FwdA, formylmethanofuran dehydrogenase subunit A.

**Supplementary Figure 7:**
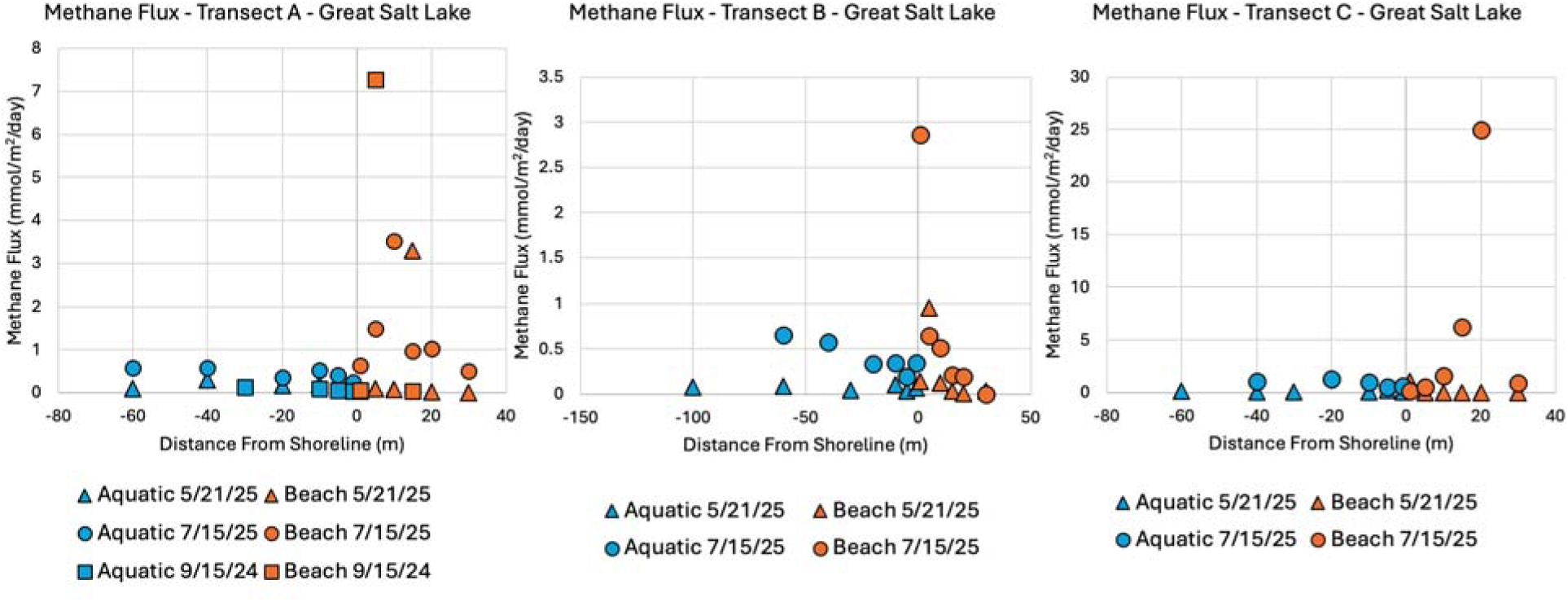
Methane flux across all dates and all transects from Bridger Bay in the GSL. The time points were selected to represent the lowest seasonal lake level (Sept. 15, 2024), the highest seasonal lake level (May 21, 2025), and the time of year which *Ca.* Methanohalophilus hillemani was collected from the lake (July 15, 2025) (see Supplementary Figure 10).

**Supplementary Figure 8:**
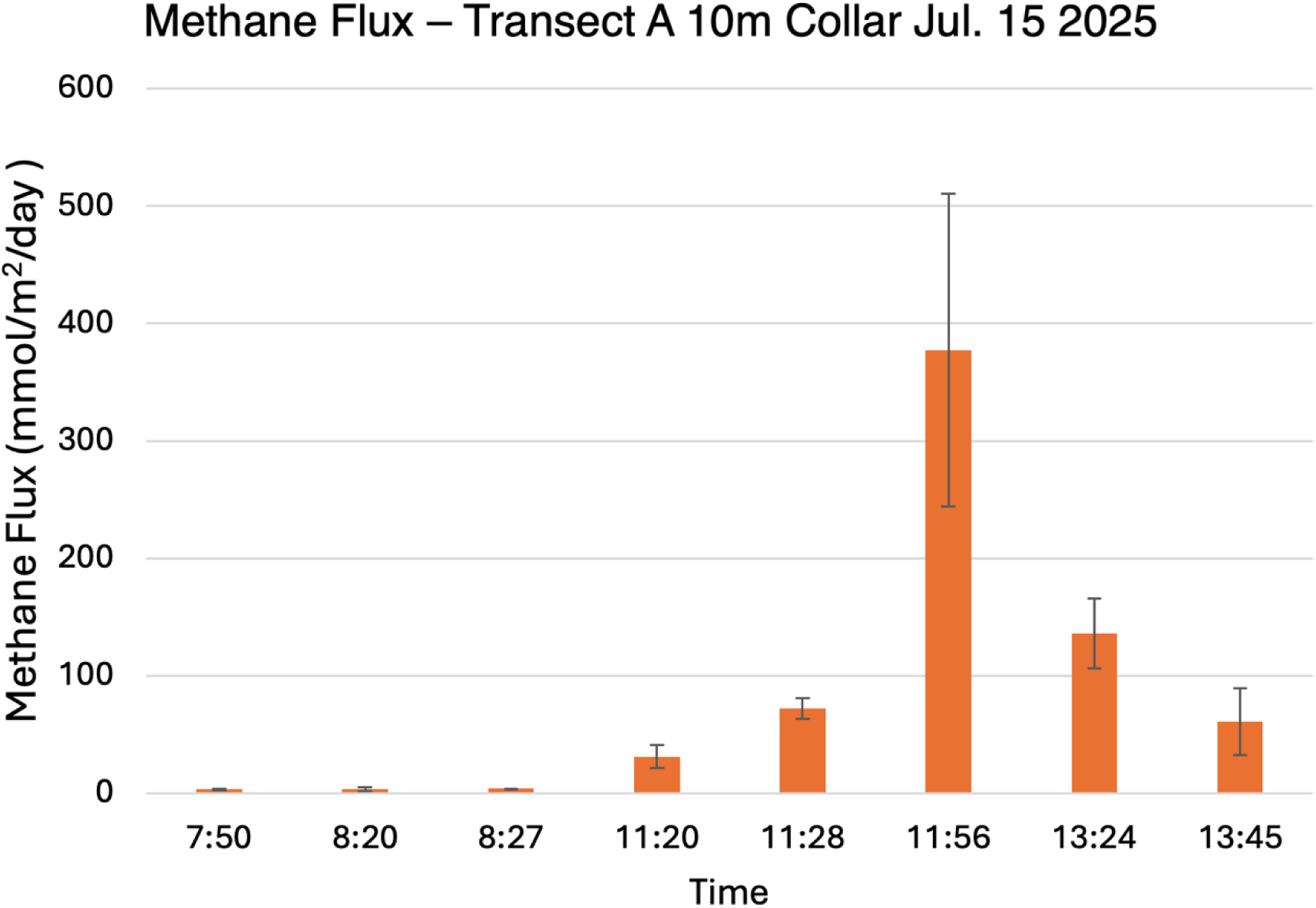
Methane flux dynamics from a single location. Terrestrial flux measurements made over the course of 6 hours from the exact same location, highlighting the variability of methane flux. This was the exact location from which the sediment core was collected that was used to generate the amplicon data presented in main text Figure 4. This was the most extreme example of flux variability observed at a single location. The atmospheric pressure was dropping that afternoon and the wind was increasing, so it is possible that a low-pressure system evacuated gas from the sediment, resulting in high methane fluxes. However, the pressure continued dropping, as did methane flux, from 11:56 to 13:45. Unless all the methane that had been “stored up” in the sediment was evacuated, meteorology cannot entirely explain the extreme variability in methane flux.

**Supplementary Figure 9:**
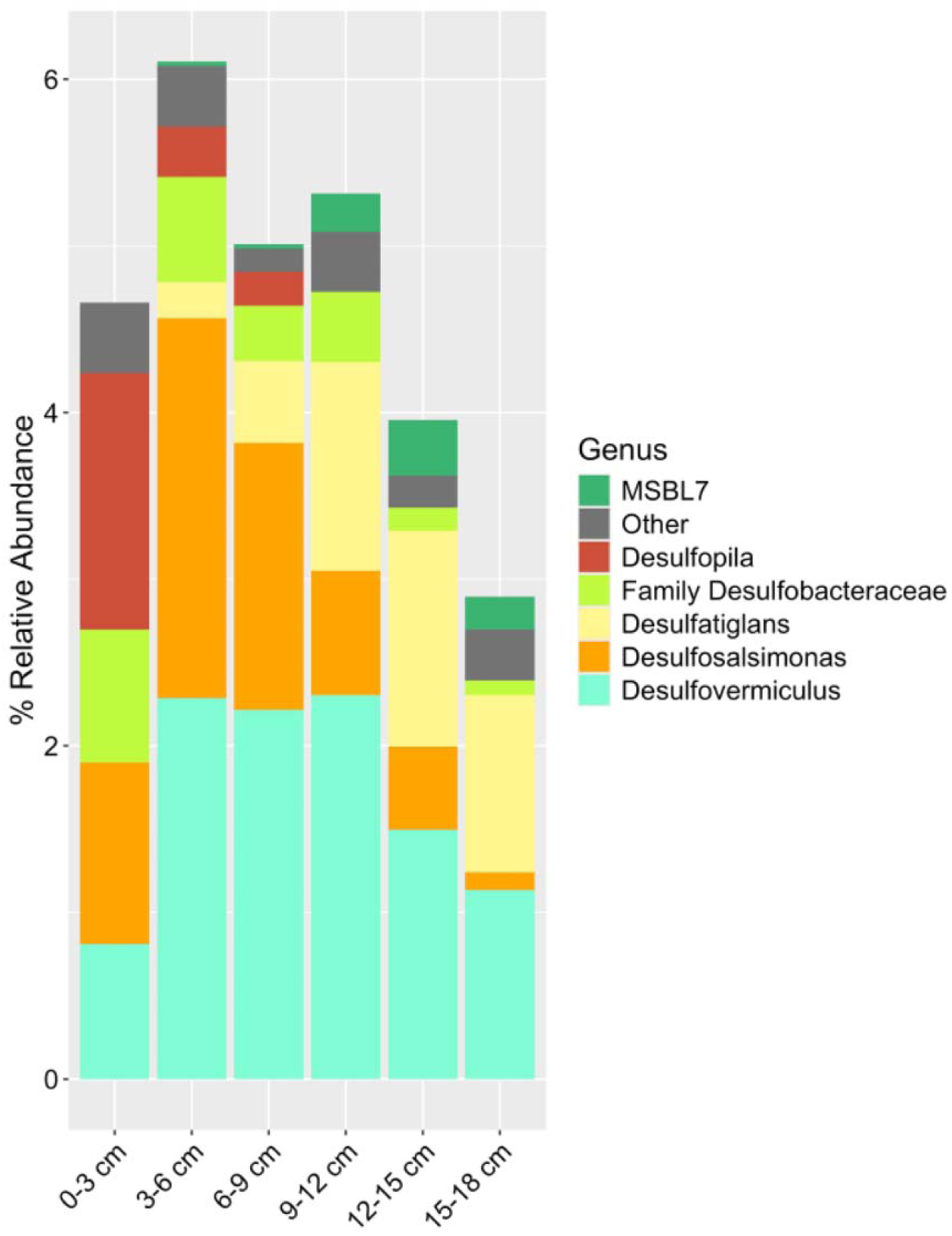
*Desulfovermiculus* is the most abundant SRB present in GSL soils. Relative abundance of the different SRB genera from a sediment core taken from the Transect A 10m flux site from July 15, 2025 based on 16s rRNA gene amplicon sequencing. Amplicons plotted belong to the taxonomic Classes *Desulfovibrionia*, *Desulfobacteria*, *Desulfobulbia*, *Desulfuromonadia*, *Desulfotomaculia*, and *Desulfitobacteriiain*, and excluded orders shown in the literature not to perform sulfate reduction. Amplicon sequence variants that constituted less than 0.1% of the total community were clustered into the “Other” category. Sequence variants that could not be assigned a genus are labeled “Family” prior to the name.

**Supplementary Figure 10:**
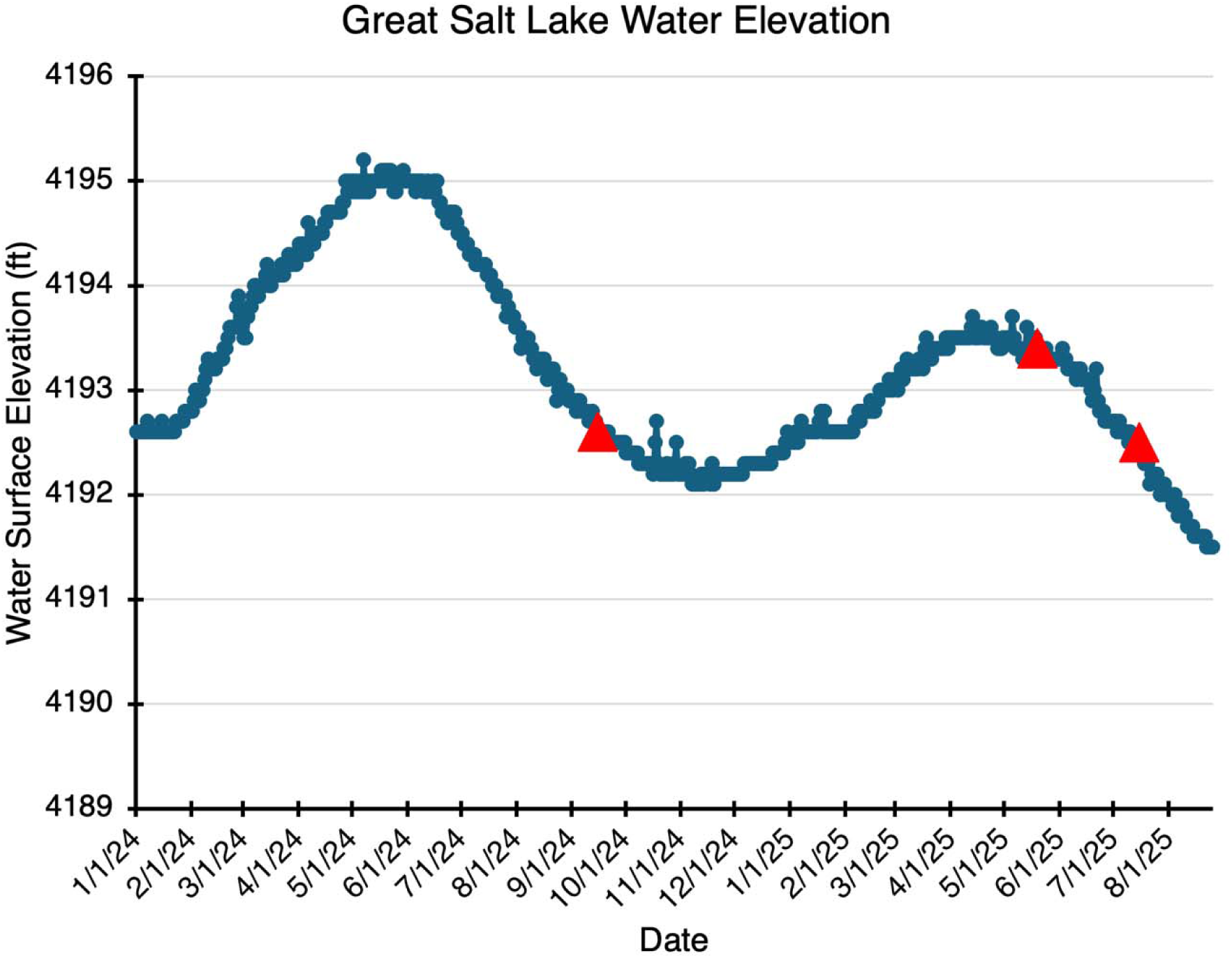
Methane flux sampling dates compared to historical lake levels. Red triangles indicate the date that methane flux measurements were taken from the GSL. Lake level data was retrieved for Saltair Boat Harbor from the USGS website.

**Supplementary Figure 11:**
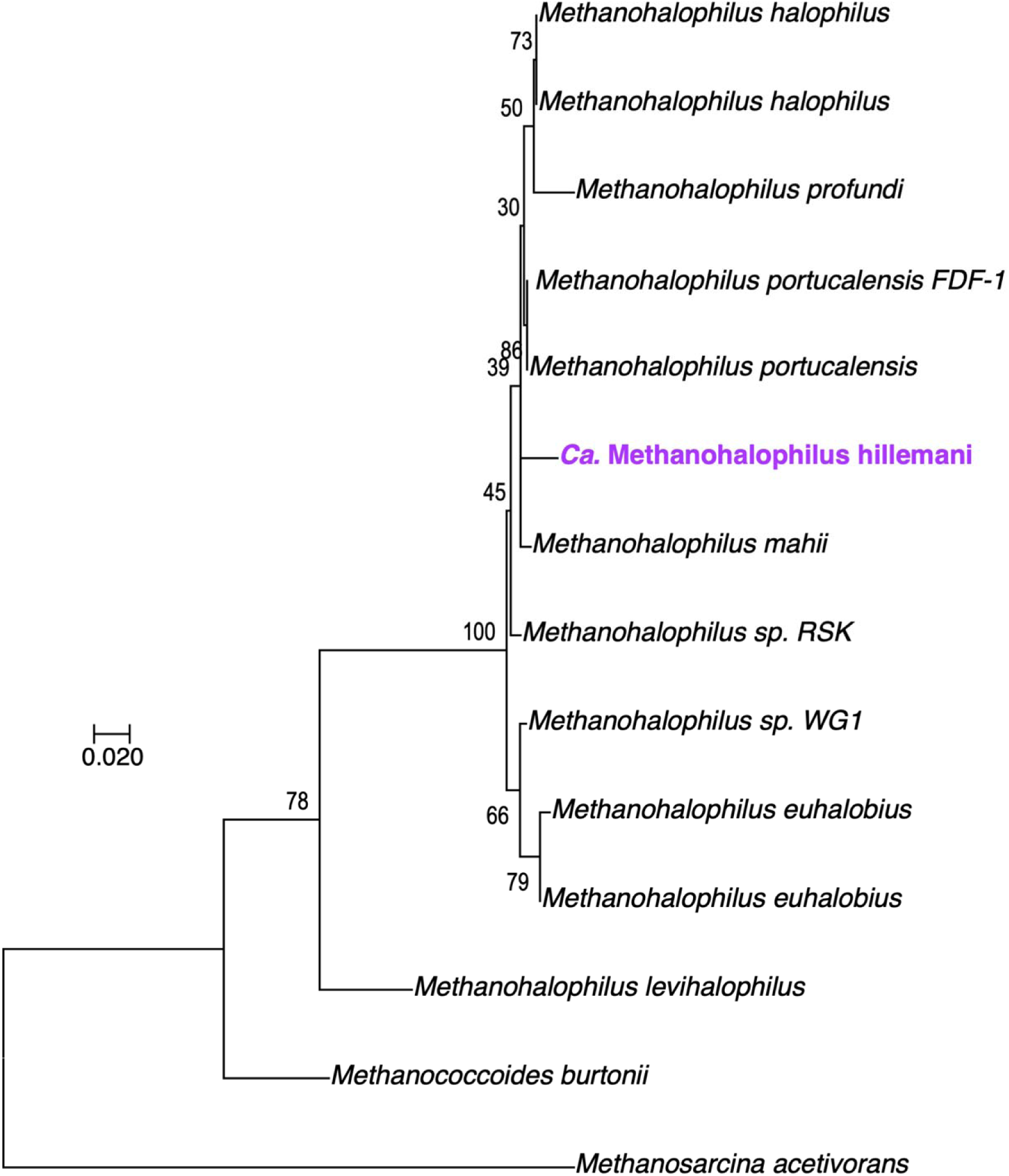
Phylogenetic tree of *Methanohalophilus* McrA sequences. Maximum likelihood tree of 573 amino acid positions constructed with IQtree2 using 1,000 ultrafast bootstrap replicates.

## Supplementary text

### Supplementary methods

#### Enrichment

Microbialite material was incubated in gas-sealed bottles with filter-sterilized GSL water collected on July 12, 2022 (salinity of 23.7%), and spiked with 10 mM MMA to enrich for methanogens [1]. From July 2022 to December 2022, enrichments were maintained by transferring inoculum, in a 1:10 ratio, into fresh lake water + MMA. When the GSL water collected in July 2022 was depleted, enrichments then used filter-sterilized GSL water collected on April 13^th^ 2023 (salinity of 16%) as the liquid media until October 2023. Lake water was then replaced by media designed by Paterek and Smith [2], originally used to isolate *Methanohalophilus mahii*, but adjusted to salinity of GSL water from April 2023. In February 2024, TMA replaced MMA as the methanogenic substrate, as it yielded more methane production. In December of 2024, we stopped adding sodium sulfide and the volatile fatty acid solution from the Paterek and Smith 1985 [2] recipe – no impact was observed on methane production. Enrichment cultures have been maintained in this manner since December 2024. More details on exact conditions and media preparation can be found in the ‘Culturing Conditions’ section.

#### DNA Extractions and 16S rRNA gene amplicon sequencing

In April of 2024, after ∼660 days of incubation and 23 transfers, phenol-chloroform-isoamyl alcohol DNA extractions were done on the methanogenic enrichments being grown in the Paterek and Smith media [2]. 1.5 mL of culture was harvested and pelleted at 20,000 x G for 10 minutes. The supernatant was discarded and the pellet was resuspended in 100 µL of 20 mM Tris-HCl (pH 8). 2 µL of 1% SDS was added to the tubes, which were then vortexed for 2 minutes. 100 µL of 25:24:1 phenol:chloroform:isoamyl alcohol was added to the tubes which were scraped against the tube rack to mix. The tubes were then centrifuged again, as before, and the top layer (∼100 µL) was pipetted into fresh tubes containing 75 µL of ice-cold isopropanol. These tubes were then vortexed, and spun at 20,000 x G for 5 minutes, the supernatant was decanted, and 500 µL of 75% ethanol was added to the (frequently invisible) pellet. The tubes were then stored at -20 °C overnight. The next day, the tubes were spun at 20,000 x G for 5 minutes and the supernatant was decanted. Tubes were then dried in a 37 °C incubator until all ethanol had evaporated off (∼20 minutes) and the (frequently invisible) pellet was resuspended in 32 µL of 20 mM Tris-HCl (pH 8).

16S rRNA gene amplification, barcoding, sequencing, and subsequent bioinformatics data analysis was performed as described in Krukenberg et al. [3]. The V4 hypervariable region of the 16S rRNA gene was amplified using the 515F and 806R Earth Microbiome Project primers, allowing approximately genus-level resolution of most previously characterized bacterial and archaeal taxa [4, 5]. The amplicon sequencing was performed via Illumina MiSeq by SeqCenter (Pittsburgh, PA). QIIME2 [6] was used to process the data and DADA2 was used to merge forward and reverse reads [7].

#### Metagenome sequencing

Metagenome libraries were constructed from the remaining DNA of a 16S rRNA amplicon sample that contained 63% relative abundance of *Methanohalophilus*. Long-read sequencing libraries were prepared using the rapid barcoding kit (SQK-RBK114-24; Oxford Nanopore Technologies) according to the manufacturer’s protocol. Sequencing was performed on a MinION device equipped with an R10.4.1 flow cell (Oxford Nanopore Technologies). Basecalling was conducted with Dorado v0.7.2 (Oxford Nanopore Technologies). Short-read sequencing libraries were generated on an Illumina NovaSeq X Plus platform (SeqCenter, Pittsburgh, PA, USA) generating 2 x 151 bp paired-end reads.

#### Metagenome assembly and annotation

Nanopore reads were assembled with MetaFlye 2.9.4-b1799 [8]. Manual evaluation of the resulting 85 contigs (≥2 kb) with SendSketch (BBTools Suite v38.94; [9]) and BLASTn (v2.14.0+) identified 10 *Methanohalophilus*-like contigs (2.21 Mb), including a 1.78 Mb fragment. These contigs were used as references for mapping Illumina short-reads with BBsplit (default parameters) and Nanopore reads with Minimap2 v2.26-r1175 [10] using default parameters prior to hybrid assembly with Unicycler v0.5.0 (--mode bold) [11]. The curated assembly produced 4 scaffolds (2.08 Mb), which were subsequently used for a second round of Illumina and Nanopore read mapping and assembly with Unicycler. Curation of this second assembly resulted in 2 *Methanohalophilus*-like contigs (2.10 Mb), which were further closed with LongStitch v1.0.5 [12] and Nanopore read mapping.

The closed genome was polished with long-reads using four rounds of Racon v1.5.0 [13] followed by one round of Medaka v2.1.0 (Oxford Nanopore Technologies) with the r104_e82_400bps_hac_v5.0.0 model. Short-read polishing was performed with two rounds of Polypolish v0.6.1 [14] and one round of POLCA v4.1.1 [15], both utilizing BWA v0.7.17-r1188 [16] alignments. The final circular genome was 2,093,478 bp in length with a GC content of 41.5 %. Genome completeness (99.99 %) was estimated using CheckM2 v1.0.1 [17].

Illumina reads were also assembled with MetaSpades v3.15.5 [18]. Coverage was calculated with BBmap (ambiguous=random), and sequences ≥2 kb were binned using MetaBAT v2.12.1 (with and without coverage; [19]), MaxBin v2.2.7 [20], CONCOCT v1.0.0 [21], Autometa v1 (archaeal and bacterial modes; including the machine learning option; [22]), and MetaDecoder v1.0.19 [23]. Final bin evaluation was performed with DAS Tool v1.1.7 [24], resulting in fragmented but complete (>99.5 %) genomes of *Halanaerobium* and *Desulfovermiculus* spp. at ∼17X and ∼25X lower coverage, respectively, compared to *Methanohalophilus* (Supplementary Table 11). Genomes were annotated with Prokka v1.14.6 [25] using both default and custom in-house annotation databases. These genomes can be found with the following accession numbers for *Ca.* M. hillemani str. BB.*, Desulfovermiculus* sp., and *Halanaerobium* sp., respectively: SAMN54119603, SAMN54119604, SAMN54119605

#### Marker gene analyses and comparative genomics

Publicly available *Methanohalophilus* spp. 16S rRNA sequences were aligned and masked using SSU-ALIGN v0.1.1 [26], resulting in a final alignment of 1,376 positions. Maximum likelihood analysis was performed with FastTree v2.1.11 [27] using default parameters. A single nucleotide polymorphism in one of the three *Ca.* M. hillemani 16S rRNA gene copies did not affect phylogenetic placement. Average 16S rRNA gene sequence identities were determined with pairwise BLASTn comparisons.

Sequence alignments of *Methanohalophilus* spp. McrA were generated with MAFFT-linsi v7.522 [28] and trimmed with trimAL v1.4.rev22 [29] using a 50 % gap threshold, producing a final alignment of 573 positions. Maximum likelihood phylogenetic analysis was conducted with IQtree2 v2.0.6 [30] using the LB+C60+F+G model and 1,000 ultrafast bootstrap replicates (Supplementary Figure 11).

A set of 65 single-copy marker proteins (Supplementary Table 12) were collected from the genomes of isolated *Methanohalophilus* spp., aligned with MUSCLE v3.8.1551 [31], and concatenated to generate a final alignment of 14,606 positions. Phylogenetic inference was performed with FastTree (option -wag) using 1,000 ultrafast bootstrap replicates.

Whole-genome comparisons were conducted to calculate average nucleotide identity (ANI) with fastANI (-fragLen 2,000; [32]) and average amino acid identity (AAI) with CompareM v0.1.2 (default parameters; https://github.com/dparks1134/CompareM), respectively. Additionally, Mauve v20150226 [33] was used to identify orthologous protein-coding sequences (≥70 % amino acid ID and ≥70 % aligned length) (Supplementary Tables 13 - 15). Orthologs were further evaluated with BLASTp alignments, and proteins were considered unique if they exhibited <50 % amino acid identity or aligned with <25 % of any other *Methanohalophilus* proteins (Supplementary Table 16, 17).

#### Culturing conditions

Since December 2024, *Ca.* M. hillemani cultures were grown in Paterek and Smith 1985 media [2], with differing concentrations of salts and TMA, and without the addition of volatile fatty acid solution and sodium sulfide. The headspace consisted of a 50/45/5 ratio of H_2_/N_2_/CO_2_ at ambient pressure (∼90kPa in Bozeman, MT). These conditions served as the control for the methane production and proteomics experiments. Cultures were grown with 10 mL of liquid medium in a 70 mL borosilicate glass serum bottle. The salt levels in our media were increased to match GSL lake water collected on Apr. 13^th^ 2023, as measured by the Joye Lab at The University of Georgia. The media was made by adding the following salts: 100 g/L NaCl, 45 g/L MgCl_2_, 5.5 g/L KCl, 1.5 g/L CaCl_2_, 0.13 g/L LiCl, 19 g/L Na_2_SO_4_, 0.5 g/L NH_4_Cl. This results in ion concentrations for Na^+^, Cl^-^, Ca^2+^, Mg^2+^, K^+^, and SO_4_ ^2-^ of 2,040, 2,770, 14, 221, 74, and 134 mM, respectively. This salt solution was then autoclaved using a 30-minute sterilization liquid cycle. After autoclaving, the media was immediately sparged for 10 minutes with 90/10 N_2_/CO_2_ gas to purge O_2_. The media bottle was then sealed and left to cool. Once cooled, the following components were added to make 1 L final volume of media: 500 µL of 1 g/L sodium resazurin solution, 10 mL of DSMZ 141 trace element solution (Modified Wolin’s mineral solution), 23.81 mL of 1 M sodium bicarbonate, 18.87 mL 1 M sodium carbonate, 5 mL of 40 g/L 2-mercaptoethane sulfonic acid sodium salt (Coenzyme M), and a small spatula tip (1-2 mg) of sodium dithionite as a reducing agent. Finally, pH was adjusted to ∼7.5. When new bottles were to be inoculated, 9.5 mL of media was added to the 70 mL serum bottles. Then, 10 µL of 50 mg/mL vancomycin and streptomycin were added to each bottle along with 200 µL of 1 M TMA and 300 µL of carbon source mix. This mix consisted of 33.33 g/L of yeast extract, 33.33 g/L of trypticase peptone, and 16.67 g/L of cysteine HCl [2]. Cultures were inoculated with 100 µL from the previous culture and then grown to stationary at 30 °C. Cultures were grown for approximately a month before being transferred to fresh media.

#### Isolation efforts

Dilution to extinction and plating were attempted to isolate *Ca.* Methanohalophilus hillemani. Solid media were prepared by adding 16 g/L agar to the medium before autoclaving. Plates were poured and streaked from active liquid cultures in a Coy chamber (Coy Lab Products, Grass Lake, MI, USA). The plates were then placed into a gas-sealed metal container (Liquid Impact Water Jet & Fabrication Fort Collins, CO, USA) and incubated at 30 °C. We tried 100% N_2_ and 50/45/5 H_2_/N_2_/CO_2_ headspaces in the gas-sealed container. Adding additional antibiotics, chloramphenicol and kanamycin to solid and liquid media, was also attempted. Dilution to extinction and plating efforts resulted in methane-producing cultures, but, based on microscopy, neither resulted in an isolate.

#### Metaproteomics sample preparation

The harvested cultures were sent to the Pacific Northwest National Laboratory (Richland, WA, USA) where protein digestion was performed using S-Trap microfilters (Protifi, Fairport NY) following an optimized procedure adapted from the manufacturer’s guidelines with reagents purchased from Sigma (St. Louis, MO, USA). Cell pellets were resuspended in 50 µL of 5 % (w/v) sodium dodecyl sulfate (SDS) prepared in 50 mM triethylammonium bicarbonate (TEAB; pH 7.5, LC-MS–grade water) and lysed by alternating brief sonication and vortex agitation until complete solubilization was achieved. Total protein concentration was quantified by bicinchoninic acid (BCA) assay (Thermo Scientific, Waltham, MA, USA), and 200 µg of protein from each sample was used for digestion. Disulfide bonds were reduced by addition of dithiothreitol (DTT) to a final concentration of 20 mM, followed by incubation at 95 °C for 10 min. After cooling to ambient temperature, cysteine residues were alkylated with 40 mM iodoacetamide in the dark for 30 min.

Samples were acidified with phosphoric acid to a final concentration of 1.2% (v/v) and diluted six-fold with S-Trap binding buffer composed of 90% methanol and 100 mM TEAB (pH 7.1). The mixtures were gently vortexed and applied to S-Trap filter cartridges. Proteins were captured on the filter by centrifugation at approximately 2,000 × g for 30–60 s, and the flow-through was reapplied twice to ensure complete binding. The filters were then washed three times with 200 µL of binding buffer to remove residual detergents and contaminants.

Bound proteins were digested on-filter with sequencing-grade trypsin (Promega, Madison, WI) and Lys-C (Fujifilm Wako, Richmond, VA) in 150 µL of 50 mM TEAB, pH 8.0 (digestion buffer) at 37 °C for 16 h. To prevent membrane desiccation during the overnight incubation, a digestion volume of at least 100 µL was maintained. Peptides were sequentially eluted using three stepwise buffers (each 80 µL): 50 mM TEAB, 0.2% formic acid in water, and 50% acetonitrile containing 0.2% formic acid. The combined eluates were dried by vacuum centrifugation and desalted using C18 macro-spin columns (Harvard Apparatus, Holliston, MA, USA). Cleaned peptide samples were subsequently dried and stored at –80 °C until LC-MS/MS analysis.

Raw proteomics data has been deposited at massIVE under accession number: MSV000100495

#### Metaproteomic MS measurements and data analysis

MS analysis was performed using an Orbitrap Exploris 480 mass spectrometer (Thermo Scientific, Waltham, MA, USA) outfitted with a homemade nano electrospray ionization interface using data-dependent acquisition (DDA) mode (QC plots in Supplementary Table 18). The ion transfer tube temperature and spray voltage were 300°C and 2.2 kV, respectively. Data were collected for 120 min following a 10 min delay after completion of sample trapping and start of gradient. FT MS spectra were acquired from 300 to 1800 m/z at a resolution of 60 K (AGC target 3e6) and up to the top 20 FT HCD MS/MS spectra were acquired in DDA mode with an isolation window of 0.7 m/z at a resolution of 30 K (AGC target 1e5) using a normalized collision energy of 30, dynamic exclusion time of 45 s, and detected charge state of an ion 2 to 6.

Peptide matching from the acquired datasets was performed using MSGF+ software [34] against proposed target meta-proteome composed from sequenced genomes (see ‘Metagenome Assembly and Annotation’ section) of novel members of *Desulfovermiculus*, *Halanaerobium*, and *Methanohalophilus* genera totaling 7,403 protein sequences. *Ca.* M. hillemani protein collection incorporated Prokka [25] predicted protein sequences with non-standard amino acid pyrrolysine (symbol O in FASTA file) and common contaminant sequences were added to the protein collection used. MSGF+ was used in target/decoy mode with 20 ppm parent ion tolerance, partial tryptic rule with static carbamidomethylation of cysteine (+57.0215) and variable modifications for the oxidation of methionine (+15.9949), deamidation of (+0.984) of asparagine and glutamine, and a methyl loss (-14.0157) for pyrrolysine (O) residues. Best matches from the MSGF+ searches were filtered at 1% FDR based on target/decoy model; this set of peptides was used in consequent quantitation analysis (Supplementary Table 19). Spectral counts (SC) and normalized spectral abundance factors (NSAF) were derived from counts of 1% false detection rate (FDR) detected peptides without consideration of peptide-protein specificity. For MASIC reports, to optimize peptide-protein specificity and protein coverage, the best protein for each peptide was guessed based on the full set of protein-peptide mappings.

Peptide abundances, integrated as area under the peptide elution peaks, were produced from raw spectra using in-house developed software MASIC [35]. MASIC peptide abundances were log2 transformed to remove skewness in the distribution of measured abundances (Supplementary Table 20). Transformed abundance values were normalized using mean central tendency method implemented in InfernoR software [36]. For each sample, *Ca.* Methanohalophilus hillemani peptide abundances were normalized to the relative abundance of *Ca.* M. hillemani by dividing the summed area under the curve (AUC) for *Ca.* M. hillemani peptides by the summed AUC for all peptides. The AUC for each *Ca.* M. hillemani peptide in each sample was then divided by this sample-specific relative abundance value (Supplementary Table 21). Peptides were then aggregated to the protein level by summing all peptide AUCs mapping to specific proteins (Supplementary Table 22). For protein level comparison, pairwise z-score analysis (standard deviation from the mean) between samples was used to extract significant differences in protein expression with significance attributed to a z-score of 2 or higher (Supplementary Tables 3 - 7). Unless otherwise indicated, all proteomic data presented in this study represents normalized protein abundances.

#### Methane flux data collection

Methane flux data was collected at Bridger Bay in the south arm of the GSL on three different dates: Sept, 15, 2024; May 21, 2025, and July 15, 2025. This is the same location where the original sample for cultivation was obtained. A Li-COR 7810 trace gas analyzer (Li-COR, Lincoln, NE, USA) was used to measure methane flux. For terrestrial flux, we used a Li-COR 8200–01S Smart Chamber placed on cylindrical, PVC soil collars (10 cm diameter). Three replicates were collected per sample. Each replicate consisted of a one-minute measurement with a 10 second deadband time, followed by a one-minute period to flush prior to the next replicate. The SoilFluxPro Software (Li-COR, Lincoln, NE, USA) was used to analyze the data, and the exponential fit was chosen to estimate methane flux.

Aquatic flux measurements were done using a custom-made floating flux chamber to take duplicate, one minute, flux measurements. The chamber consisted of a gas-tight coffee can fitted with a foam floatation collar. The internal volume of the flux chamber was 2,584 mL. The chamber was removed from the water’s surface between duplicates to allow the chamber gas composition to return to atmospheric conditions. Aquatic fluxes were calculated from the linear change in methane concentration over time, as previously described [37].

On each sampling date, flux was collected in a transect, perpendicular to the GSL Bridger Bay shoreline. Measurements extended toward to shore and out into the water, perpendicularly, from the shore.

#### Soil core collection and *mcrA* gene analysis

Soil cores were collected from within the boundary of the terrestrial collar with the highest methane flux (transect A - 15m for May 2025 and transect A – 10m for July 2025) and transported back to Montana State University on ice. In the lab, the core was separated into sections of three or six cm each. DNA from each soil core section was extracted with MP Bio Soil Extraction kits, and 16S rRNA gene amplicon sequencing was performed, as described above. For the July 15, 2025 core, the DNA extract was also used for amplifying and sequencing an amplicon for the gene encoding the alpha-subunit of methyl-coenzyme M reductase (*mcrA*). The mlas-mod-F and *mcrA*-rev-R primers and the touchdown amplification program from Angel et al. 2012 were utilized [38]. Sequencing and analysis of data was performed as described above. However, due to variable lengths of the *mcrA* gene amplicon, the merging of forward and reverse reads via DADA2 was not uniformly successful. Therefore, the *mcrA* gene amplicon data was analyzed using only the ∼227 bases from the forward read.

## Supplementary results

### Unique genetic features of *Ca.* Methanohalophilus hillemani

We identified 64 coding sequences in *Ca.* M. hillemani that have no homologs in any of the other 13 *Methanohalophilus* genomes (Supplementary Tables 16, 17). When analyzing these unique (among *Methanohalophilus*) sequences, there were many proteins that appeared to be part of potential operons. Three operons of unique *Ca.* M. hillemani genes involved in immune response were found. One consists of three genes encoding type 1 restriction modification system proteins. Type 1 restriction-modification systems are responsible for recognizing self-DNA (hsdS), modifying self-DNA via methylation (hsdM), and destroying foreign DNA via restriction endonuclease activity (hsdR) [39, 40]. *Ca.* M. hillemani encodes a second type 1 restriction-modification system operon, however, these proteins have significant homology to sequences from other *Methanohalophilus* species. A second unique operon encodes a toxin production protein, a CRISPR associated protein, and two antiviral radical SAM proteins. A third consists of three antibiotic resistance ABC transports and a protease. Borton et al. in 2018 suggested that viral predation may drive strain differentiation in *Methanohalophilus* species [41]. Our data corroborates that immune response is a differentiating factor between *Ca.* M. hillemani and other *Methanohalophilus* genomes.

A large operon of unique coding sequences for DNA maintenance and modification proteins was also found in the *Ca.* M. hillemani genome. This putative operon includes genes coding for a chromosome segregation protein, a DNA helicase, a DNA sulfur modification protein, and a RecB family nuclease. In addition to this operon, *Ca.* M. hillemani encodes three unique DNA helicases and a unique double strand break repair protein. DNA repair mechanisms are shared across the genus, including the DNA repair proteins RadA, B, and C. However, the unique *Ca.* M. hillemani sequences in this category indicate that DNA maintenance may be of particular importance.

Cobalamin (vitamin B12) is a cobalt-containing corrinoid cofactor that is essential for methyl-dismutating methanogenesis [42], and all *Methanohalophilus* species share most genes required for its biosynthesis. While *Ca.* M. hillemani encodes cobalamin biosynthesis, its genome also contains two operons with unique coding sequences involved in cobalamin import. The first operon encodes the Cobalamin import permease, BtuC, the Cobalamin import ATP-binding protein, BtuD, and an ABC transporter. The other operon contains a second copy of BtuD and two ABC transporters. Other *Methanohalophilus* species encode genes annotated as BtuC and BtuD; however, *Ca.* M. hillemani appears to have acquired/evolved divergent uptake pathways, as its BtuC and D proteins have low (<50%) amino acid identity with any proteins in the other genomes.

### Etymology

The newly cultured archaeon is a methanogen affiliated with the genus Methanohalophilus, family Methanosarcinaceae, order Methanosarcinales, class Methanomicrobia, phylum Methanobacteriota, for which we propose the name *Candidatus* Methanohalophilus hillemani sp. nov. strain BB.

Me. tha. no. ha. lo. phi’lus. M. L. n. *methanum*, methane; Gr. n. *halo*, salt; Gr. no. *philos*, lover; M. L. n. *Methanohalophilus*. methane-producing salt lover.

hil.le.ma’ni N.L. gen. masc. n. *hillemani,* of Hilleman; named in honor of Maurice Hilleman (1919-2005), a microbiologist and vaccinologist born in Miles City, Montana. Hilleman graduated from Montana State University and, over the course of his career, developed over 40 vaccines including eight that are currently recommended for children and adults in the United States. His vaccines significantly reduced morbidity and mortality during two 20^th^ century influenza pandemics, and his cumulative contributions are credited with saving 10s of millions of lives worldwide. See https://hillemanfilm.com/dr-hilleman for more information. Appropriately, the results of this study show that *Ca.* M. hillemani invests heavily in immune response.

The strain name BB refers to Bridger Bay on the Great Salt Lake from which *Candidatus* Methanohalophilus hillmani was cultured from.

